# scLM: automatic detection of consensus gene clusters across multiple single-cell datasets

**DOI:** 10.1101/2020.04.22.055822

**Authors:** Qianqian Song, Jing Su, Lance D. Miller, Wei Zhang

## Abstract

In gene expression profiling studies, including single-cell RNA-seq (scRNAseq) analyses, the identification and characterization of co-expressed genes provides critical information on cell identity and function. Gene co-expression clustering in scRNA-seq data presents certain challenges. We show that commonly used methods for single cell data are not capable of identifying co-expressed genes accurately, and produce results that substantially limit biological expectations of co-expressed genes. Herein, we present scLM, a gene co-clustering algorithm tailored to single cell data that performs well at detecting gene clusters with significant biologic context. scLM can simultaneously cluster multiple single-cell datasets, i.e. consensus clustering, enabling users to leverage single cell data from multiple sources for novel comparative analysis. scLM takes raw count data as input and preserves biological variations without being influenced by batch effects from multiple datasets. Results from both simulation data and experimental data demonstrate that scLM outperforms the existing methods with considerably improved accuracy. To illustrate the biological insights of scLM, we apply it to our in-house and public experimental scRNA-seq datasets. scLM identifies novel functional gene modules and refines cell states, which facilitates mechanism discovery and understanding of complex biosystems such as cancers. A user-friendly R package with all the key features of the scLM method is available at https://github.com/QSong-WF/scLM.

## Introduction

Co-expressed genes work in concert in biological pathways and processes [1–3]. Such genes are involved in crucial biological activities including immune cell activation [4, 5], cellular epithelial-mesenchymal transition (EMT) [6], and transcription factor-mediated gene regulatory networks and signaling pathways [7, 8]. Co-expression of genes based on similarities among their expression profiles, has been a primary way to unravel gene-gene relationships and facilitate functional annotation. Therefore, identification of co-expressed genes provides functional insights into underlying cellular and molecular mechanisms in normal and disease processes.

The recently developed single-cell RNA-sequencing (scRNA-seq) technology provides high-resolution of gene expression at the single cell level, yet presents certain challenges for gene expression analysis. In contrast to bulk RNA-seq, single-cell data has been shown to exhibit a characteristic negative binomial distribution pattern [9–12], wherein genes suffer from stochastic dropouts and over-dispersion problems. Dropouts, or genes that exhibit excessive zero values [13–15], represent a special type of missing value, which can be caused by low RNA input or stochastic expression fluctuation at the single-cell level. Over-dispersion relates to the substantially large cell-to-cell variability in gene expression profiles which likely arises from technical noise stemming from low input RNA and PCR amplification bias [16].

Rapid advances in scRNA-seq technologies have made it feasible to perform population-scale studies in which the transcriptome is measured for thousands of single cells from multiple samples or conditions [17–21]. This in turn has amplified the need for versatile gene co-expression approaches that not only address the unique challenges of scRNA-seq data, but also the challenges of datasets integration including batch effects, technical variation (e.g., mRNA quality, pre-amplification efficiency), and extrinsic biological variability.

Classical methods designed for analysis of bulk transcriptome data such as WGCNA [22] and clust [23] are not designed to account for the unique characteristics of scRNA-seq data. Some network-based approaches for single cell data, including SCENIC [24], Cell Specific Network (CSN) [25], and LTMG [26], could detect gene co-expression modules as part of the network reconstruction. However, these methods didn’t account for the technical noise and extrinsic variance among multiple samples. Therefore, there is a clear need to develop a tailored and effective method for scRNA-seq data to extract “consensus” co-expressed genes, that is, to extract the genes that are consistently co-expressed in each of the multiple datasets.

Herein, we developed a novel method, single-cell Latent-variable Model (scLM), to simultaneously extract co-expressed genes that exhibit consensus behaviors from multiple single-cell datasets. The scLM method accounts for both cell-level covariates and sample-level batch effects. We assessed the performance of scLM in both simulated data and experimental data. The scLM achieved the best performance over other commonly used methods. We then applied the scLM to our in-house scRNA-seq data generated from 4 non-small cell lung cancer (NSCLC) tumor tissues and their corresponding adjacent normal tissues. The scLM method identified tumor-specific co-expressed gene modules with significant prognostic values. Furthermore, these co-expression modules contributed to the subtle characterization of lung tumor cell states. In addition, we applied scLM to analyze a set of malignant cells from NSCLC, head and neck squamous cell carcinoma (HNSCC), and melanoma. We identified a common co-expressed gene program across different cancer types, providing insights into fundamental mechanisms of carcinogenesis.

## Methods

### Single-cell Latent-variable Model (scLM)

We propose a latent variable model to explicitly disentangle different sources of variability in population-scale scRNA-seq data. Our goal is to perform simultaneous detection of co-expressed genes across multiple single-cell conditions/datasets. Specifically, let *x*_*ijk*_ denote the gene expression level experimentally measured for the *i*’th gene (*i* ε {1,…, *m*}) in the *j*’th cell (*j* ε {1,…, *n*_*k*_}) in condition/dataset *k* (*k* ε {1,…, *K*}).

As multiple recent studies [9–12] show that the expression of most genes in single cell data is sufficiently captured by Negative Binomial (*NB*) distribution, *NB* model is chosen as an appropriate model to formulate single cell data. It is supported by the physical modeling of bursting gene expression [12, 27] and is also commonly used in scRNA-seq analysis [9–12]. Therefore, without loss of generality, we assume that the measured gene expression *x*_*ijk*_ for cell *j* in dataset *k* follows the *NB* distribution *NB*(*p*, *γ*), which has the probability function as:

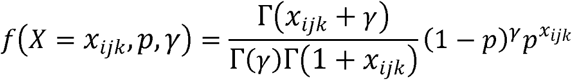

If *µ*, *θ*, and *σ*^2^ represent the mean, dispersion, and variance of this NB distribution, then we have

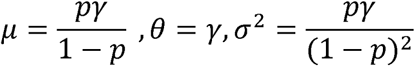

also,

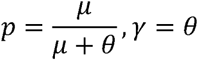

Therefore, the probability function converts to:

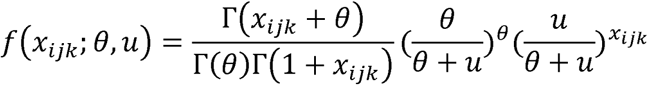

As *u* and *θ* are different regarding different genes (*i* ε {1,…, *m*}) and batches (*k* ε {1,…, *K*}), we have

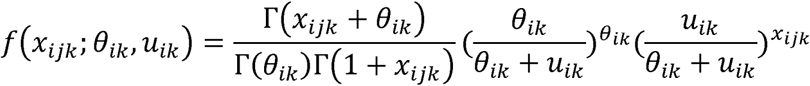

Herein, *u*_*ik*_ represents the estimation for the intrinsic gene expression level across all cells in sample *k*, *θ*_*ik*_ is the dispersion parameter, and 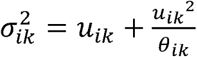 represents the square deviation of the observed gene expression level across cells in this sample.

Let **z***i* = (**z**_*i*l_, …, **z**_*i*λ_)^′^ be a vector consisting of λ unobserved latent variables that are shared by *K* different conditions/datasets. We assume the generalized linear model below

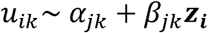

which is used to distinguish the intrinsic biological variability **z**_*i*_ from the extrinsic signals (*α*_*jk*_ and *β*_*jk*_) including the technical variances at the cell-level (j) and batch effects at the sample-level (k).That is, the *u*_*ik*_ is composed of the intrinsic biological pattern of gene *i* captured by latent variables *z*_*i*_ regardless of the confounding variabilities at the cell-level and sample-level, while variances due to technical biases and batch effects are captured by offsets *α*_*jk*_ and scale factors *β*_*jk*_ . Since **z**_*i*_ is the same for specific gene *i*, and *u*_*ik*_ is estimated from observed counts *x*_*ijk*_, we further turned the formulas into

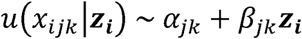

To alleviate the impact of extreme values, we utilized logarithm form in the linear model that has been frequently used [28–32] in single cell RNA-seq data, i.e. the Generalized Linear Model (GLM),

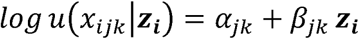

where *u*(*x*_*ijk*_ | **z**_*i*_) is the conditional mean of *x*_*ijk*_ given **z**_*i*_. In this way, the original gene expression data are projected into a λ-dimensional latent space z by the generalized linear model, with the technical biases and batch effects removed during the projection. In this latent space, the expression level of gene *i* is represented as **z**_*i*_. Since genes sharing similar expressions are located close to each other, a group of co-expressed genes will form a cluster in the latent space. Thus, different groups of co-expressed gene modules can be identified through clustering of the latent variables (Figure 1).

**Figure 1.**
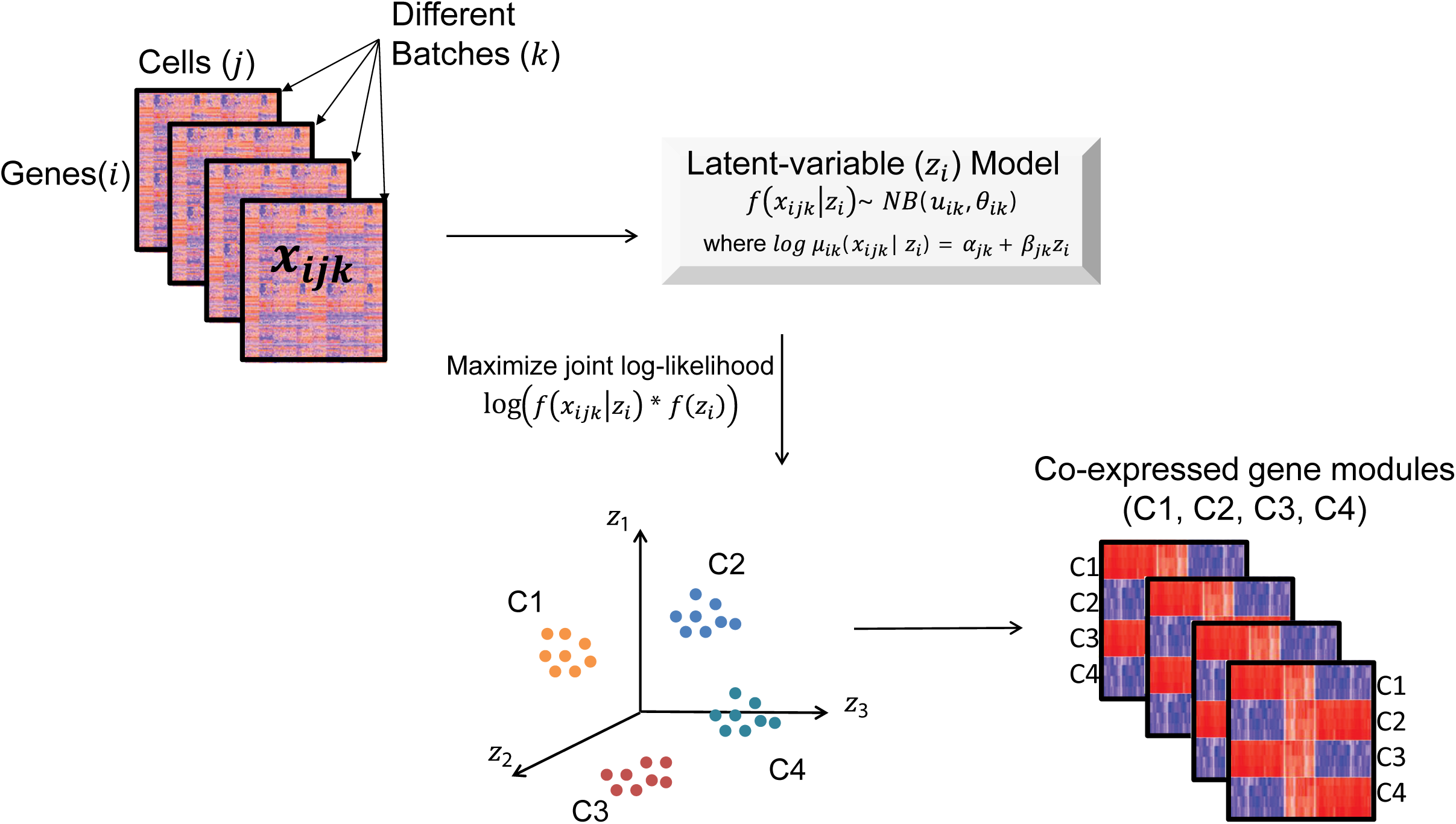
Schematics of scLM for identifying consensus co-expressed gene clusters across multiple datasets. Schematic representation of how consensus co-expressed genes across multiple datasets/conditions can be discovered by the scLM method. The gene expression profiles of individual cells are disentangled by the latent variables representing the intrinsic biological signals, and the related coefficients reflecting the technical variances.

To estimate the parameters in our model, we used the maximum likelihood approach. As assumed above that *x*_*ijk*_ follows the *NB* distribution, the conditional log-likelihood function of *x*_*ijk*_ can be written as:

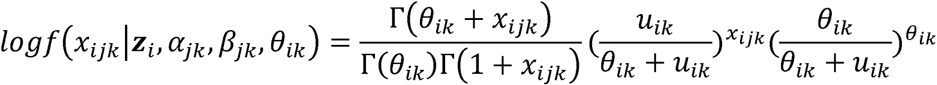

in which,

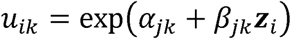

For the latent variable **z**_*i*_, *f*(**z**_*i*_) represents the density function of the standard multivariate normal distribution N(0, ***I***_λ_). Therefore, the joint log-likelihood of (*x*_*ijk*_, **z**_*i*_) can be written as

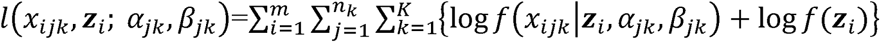

To control model complexity and overfitting, we used the least absolute shrinkage and selection operator (LASSO, L1-norm penalty), which is applied to the following penalized joint log-likelihood estimation:

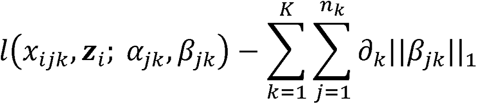

Then the above parameters are estimated by maximizing the penalized joint log-likelihood function, that is, maximizing the following penalized joint log-likelihood,

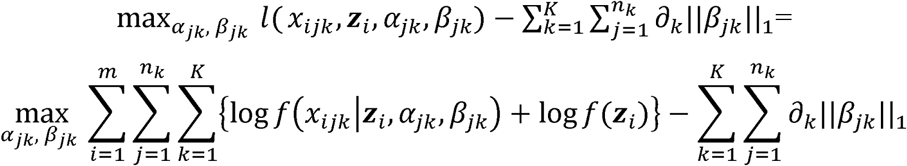

where the summation is due to the conditional independence assumption of *x*_*ijk*_ given **z**_*i*_.

To estimate the parameters *α*_*jk*_ and *β*_*jk*_, we solve the following optimization problem conditional on **z**_*i*_,

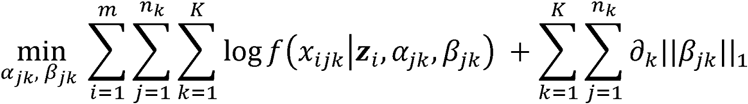

here we use the coordinate descent algorithm provided in [33], therefore optimize the above log-likelihood function. Herein the update of parameters *α*_*jk*_ and *β*_*jk*_ depends on **z**_*i*_.

As the latent variables **z**_*i*_ are not observable in our model, we use the Markov Chain Monte Carlo (MCMC) simulation to iteratively update *z*_*i*_, for maximizing the penalized joint log-likelihood. That is, we replace the value in the parameter updates by its expectation with respect to **z**_*i*_, through repeatedly sampling the latent variable **z**_*i*_ from the following joint posterior distribution, i.e.

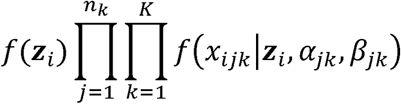

With the estimated latent variables **z**_*i*_, that is, with the genes projected into the latent space, we cluster genes that are projected in the latent space to identify co-expressed genes. Here we use K-means clustering to divide genes into λ clusters based on the latent variables **z**_*i*_. The parameter λ can be either determined according to the Bayesian information criterion (BIC), or chosen by user’s preference. All source codes of the scLM software is open released in the GitHub repository: https://github.com/QSong-WF/scLM.

### Data generation in simulation studies

Based on the single cell data characteristics, we used the negative binomial distribution to simulate two synthetic cohorts (SD1, SD2). Each synthetic cohort contains 9 sets of simulated gene expression data with an increasing number of datasets (D1 – D9). That is, D1 contained one individual dataset (n = 1), D2 contained two individual datasets (n = 2), …, and D9 contained nine individual datasets (n = 9). Each individual dataset contains 180 genes belonging to three clusters, with 60 co-expressed genes in each of the three clusters. For each gene cluster *c* ε {1,2,3} in batch *n*, their gene expression was sampled from the negative binomial *NB*(*u*_*cn*_, *θ*_*cn*_) distribution, where *u*_*cn*_ and *θ*_*cn*_ refer to the mean and deviation. Different gene clusters have different values of *u*_*cn*_ and *θ*_*cn*_. Full expression values and cluster membership for these datasets are provided in the scLM example data.

Additionally, we utilized the Splatter package [34] to generate another two batches of simulated data with dropout effects, which more accurately recapitulate actual scRNA-seq data distributions. Specifically, we adjusted the batch parameter ‘batch.facLoc’ and ‘batch.facScale’ as 1 and generated 16 different batches of data. Each batch consists of 240 cells, and 240 genes constituting four groups of co-expressed genes as the group truth, which was achieved by adjusting the ‘de.prob’ parameter. We also added the dropout effects in these simulation data by setting ‘experiment’ for global dropout and the ‘dropout.mid’ parameter as 0.1. These 16 batches of data made up the synthetic dataset SD3 and SD4. Full expression values are provided in the scLM example data.

### In-house and public single-cell data

#### In-house dataset

Fresh tumor and adjacent normal tissues from four NSCLC patients were collected by the Tumor Tissue and Pathology Shared Resource (TTPSR) into Miltenyi Tissue Storage Medium (130-100-008; San Diego, CA). Tissues were then processed to single-cell suspensions using the Miltenyi human Tumor Dissociation Kit (130-095-929) and the gentleMACS Octo Dissociator with Heaters (130-096-427). Red blood cells were removed by negative selection using Miltenyi CD235a (Glycophorin A) microbeads (130-050-501) and LS Columns (130-042-401). Recovered cell numbers were determined by trypan blue exclusion using a LUNA II automated cell counter (L40001; Logos Biosystems, Inc.; Annandale, VA). In preparation for scRNAseq, cells were thawed and washed according to the demonstrated protocol developed for human PBMCs by 10X Genomics (San Francisco, CA).

All scRNAseq procedures were performed by the Cancer Genomics Shared Resource (CGSR) of the WFBMC-CCC. Viable cells in suspensions were loaded into wells of a 10X Genomics Chromium Single Cell A Chip (Kit PN-120236). Single-cell gel beads in emulsion (GEMs) were created on a Chromium Single Cell Controller and scRNA-seq libraries were prepared using the Chromium Single Cell 3′ Library and Gel Bead kit v2 (PN-120237). Sequencing libraries were loaded at 1.3 PM on an Illumina NextSeq500 with a High Output 150 cycle kit (FC-404-2002; Illumina, San Diego, CA) for paired-end sequencing. A total of 11,813 single cells were captured, with the number of cells recovered per channel ranging from 369 to 2,502. Low-quality cells were discarded if the cell number with expressed genes was smaller than 200. Only malignant cells from four tumor samples and epithelial cells from three adjacent normal samples were used in this study. The scRNA-seq data were deposited in the GEO of NCBI database (GEO Accession: GSE117570) at https://onlinelibrary.wiley.com/doi/full/10.1002/cam4.2113 [35].

#### Melanoma dataset

We downloaded the expression matrix data of melanoma from the GEO of NCBI database (GEO Accession: GSE72056) at https://www.ncbi.nlm.nih.gov/pubmed/27124452 [36]. This dataset includes expression profiles of 23,689 genes in 4,645 cells from 19 melanoma tumors. These cells include both malignant cancer cells and non-malignant cells. For our analyses, samples with more than 200 malignant cells and genes expressed in over 300 cells were included in the input matrix.

#### Head and neck cancer dataset

We downloaded the expression matrix data of the head and neck cancer dataset from the GEO of NCBI database (GEO Accession: GSE103322) at https://www.sciencedirect.com/science/article/pii/S0092867417312709 [6]. This dataset consists of 5,902 cells from 18 patient samples after initial quality controls, including 2,215 malignant and 3,363 non-malignant cells. For our analyses, we used the samples with more than 200 malignant cells and genes expressed in over 300 cells as the input matrix.

## Breast cancer dataset

We downloaded the expression matrix data of breast cancer single-cell RNA-seq dataset from the GEO of NCBI database (GEO Accession: GSE118390) at https://www.nature.com/articles/s41467-018-06052-0 [17]. For our analysis, we used malignant cells and genes expressed in over 300 cells as input.

## Clustering evaluation index

Each clustering result produced by applying a specific clustering method to a specific dataset was assessed using the following cluster evaluation indices: the Adjusted Rand Index (ARI) [37], the Calinski-Harabasz (CH) index [38], the Davies-Bouldin (DB) index [39], and the Dunn index [40]. CH index evaluates the cluster validity based on the average between- and within-cluster sum of squares. DB index is obtained by averaging all the cluster similarities. The smaller the index is, the better the clustering result is. Dunn index uses the minimum pairwise distance between objects in different clusters as the inter-cluster separation and the maximum diameter among all clusters as the intra-cluster compactness.

## Cell clustering based on co-expressed gene modules

With the co-expressed gene modules, we utilized mean value of the modules in each single cell as input for graph-based clustering, and UMAP for visualizing cell clusters. Graph-based clustering was performed using the Seurat package (v3.1), while UMAP analysis was performed using the umap package (v.0.2.3.1) [41] in R (v.3.4.3). The number of epochs (n_epochs) was set at 20. The n_neighbors value was set at 15, and min_dist was set as 0.1.

## Statistical analysis

Kaplan-Meier (KM) analysis was performed using the ‘survival’ R package (http://cran.r-project.org/web/packages/survival/index.html). Log-rank test was used to test the differences of survival curves. t-test is used to calculate P-value when evaluating the outperformance of scLM.

## Functional Analysis

### Hallmark collection

We downloaded the Hallmark gene set collection for functional analyses from the Molecular Signatures Database (MSigDB) [42], a widely used and comprehensive database. Each hallmark in this collection consists of a “refined” gene set that conveys a specific biological state or process and displays coherent expression. The hallmarks effectively summarize most of the relevant information of the original founder sets and, by reducing both variation and redundancy, provide more refined and concise inputs for gene set enrichment analysis.

### Pathway database

Reactome (http://www.reactome.org) is a manually curated open-data resource of human pathways and reactions, which is an archive of biological processes and a tool for discovering potential functions. Gene sets derived from the Reactome [43] and KEGG [44] pathway database were downloaded from the MSigDB Collections.

### Enrichment test

Functional enrichment based on the Reactome and GO terms databases was assessed by hypergeometric tests, which were used to identify *a priori*-defined gene sets that showed statistically significant differences between two given clusters. The test was performed by the clusterProfiler package [45]. We further corrected the test P-values by the Benjamini-Hochberg correction; P-values less than 0.05 were considered statistically significant.

## Results and discussion

### Overview of the scLM

We developed a new method, single-cell Latent-variable Model, scLM, for simultaneously identifying consensus co-expressed genes from multiple scRNA-seq datasets. Our hypothesis is that co-expressed genes coordinating biological processes can be captured across multiple different datasets. In our model, we assumed that latent variables could capture the intrinsic signals of the co-expressed genes regardless of technical variances and batch effects among different datasets. Figure 1 provides an illustrative overview of the scLM method. Briefly, the input contains a collection of multiple datasets (*k*) representing the single-cell sequencing data generated under different clinical or experimental settings. In the *k*-th dataset, we assume that the observed expression levels, *x*_*ijk*_, of the *i*-th gene across cells *j* ε {1, …, *n*_*k*_) follow the negative binomial distribution *NB*(*µ*_*ik*_, *θ*_*ik*_). The intrinsic biological variability of gene i across all cells and all datasets is captured by the latent variables *z*_*i*_ in a λ-dimension latent space. This is achieved through a conditional generalized linear model (GLM) log *u*(*x*_*ijk*_ | *z*_*i*_) - *α*_*jk*_ + *β*_*jk*_ *z*_*i*_ that distinguishes the intrinsic biological variability *z*_*i*_ from the extrinsic signals (*α*_*jk*_ and *β*_*jk*_) including the technical variances at the cell-level (*j*) and batch effects at the sample-level (*k*). The latent variables and other parameters are estimated and obtained using Markov Chain Monte Carlo (MCMC) and maximum likelihood approaches. Therefore, different groups of co-expressed genes (C1-C4) across multiple datasets can be identified through clustering genes in the latent space. Further explanations of the mathematical model representations are included in Methods. The software to implement the scLM method is available at https://github.com/QSong-WF/scLM.

### Performance evaluation on simulation data

To evaluate the performance of scLM, we benchmarked it against other methods, including LTMG [26], SCN [25], Seurat-wgcna, MNN-wgcna, and SCENIC [24]. Seurat-wgcna and MNN-wgcna refer to the co-expression analysis using WGCNA [22], following the batch correction by Seurat [46] or MNN [47]. As SCENIC relies on the RcisTarget database that required real gene input, we omit comparing with SCENIC on simulation data but still include it in the comparison on real single cell data.

We first generated two synthetic data cohorts (SD1, SD2) from negative binomial distribution. Each cohort contained 9 sets (D1-D9) of simulated data with an increasing number of samples. That is, D1 contained one individual dataset (n=1), D2 contained two individuals of datasets (n=2), and so on. Each set contained three co-expressed gene clusters as ground truth. Additionally, we utilized the Splatter package [34] to generate another two batches of simulated data (SD3, SD4) with dropout effects, which can more accurately recapitulate actual scRNA-seq data distributions. Details of the simulation datasets are provided in the Methods section.

With the simulated data cohorts, we applied scLM and other methods (LTMG, SCN, Seurat-wgcna, and MNN-wgcna) to identify the co-expression clusters. To assess and quantify clustering accuracy, we used the adjusted Rand index (ARI) as performance metric to rank these methods (Figure 2 A – D). The corresponding barplots represent the ARI of the identified clusters by each method compared to the ground truth. Notably, scLM accurately identified each gene cluster in four cohorts, and demonstrated much higher ARIs [37] (mean ± SE: 0.979 ± 0.063 for SD1, 0.971 ± 0.031 for SD2, 0.899 ± 0.043 for SD3, 0.886 ± 0.025 for SD4). The other methods showed relatively lower ARIs. For example, Seurat-wgcna and CSN showed lower ARI in SD1 (mean ± SE: 0.627 ± 0.028) and SD3 (mean ± SE: 0.520 ± 0.070) respectively. LTMG presented with a little higher ARI and lower variance in four data cohorts. These results demonstrate the outperformance of scLM in identifying accurate co-expressed genes from multiple datasets.

**Figure 2.**
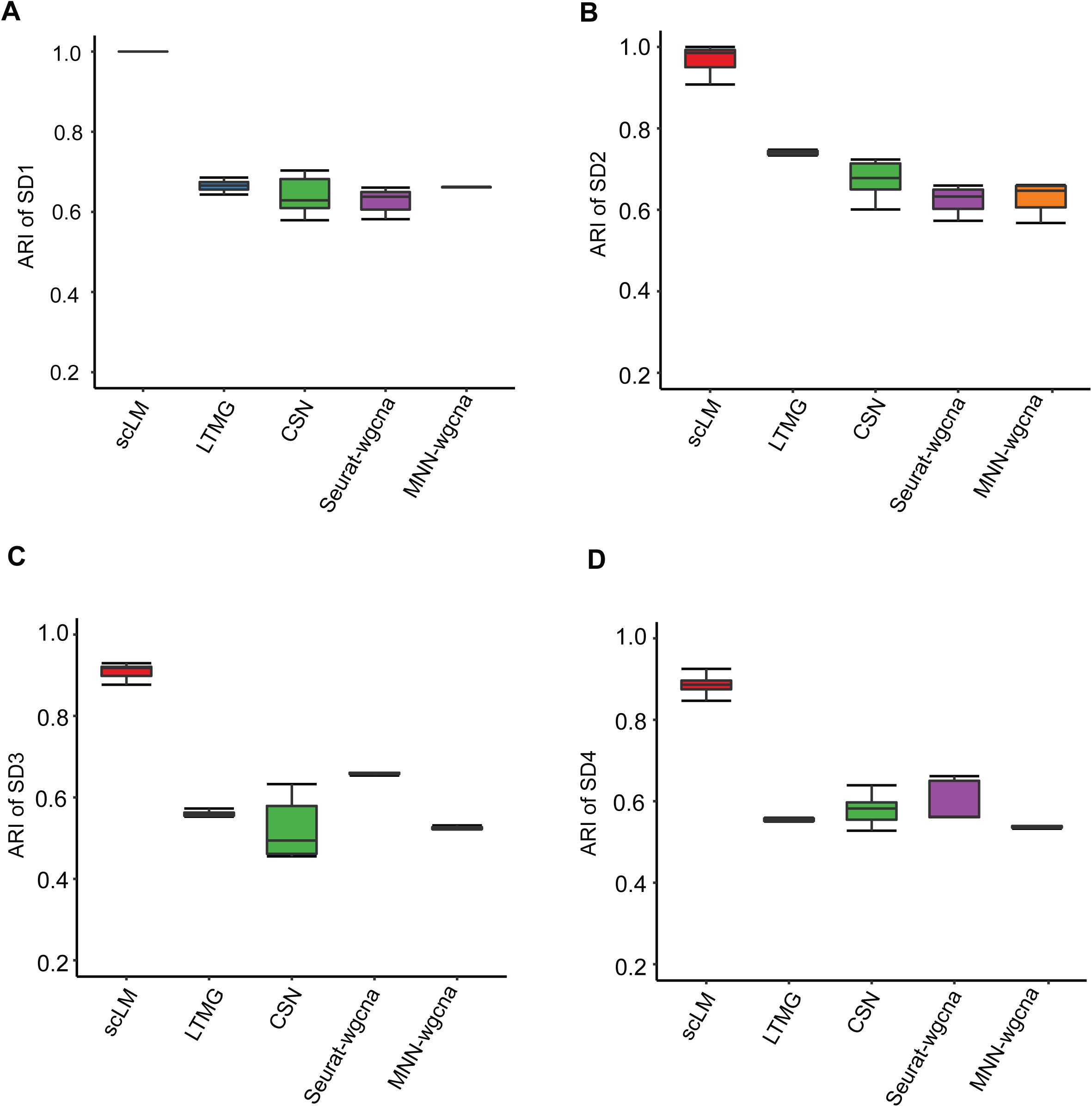
Performance evaluation on simulation data. scLM compared with the other methods (LTMG, CSN, Seurat-wgcna, and MNN-wgcna) on four synthetic cohorts (SD1 - SD4). Each synthetic cohort contains an increasing number of simulation data. The barplot represents the ARI of identified gene clusters compared to the ground truth. **(A)** ARI of SD1; **(B)** ARI of SD2; **(C)** ARI of SD3; **(D)** ARI of SD4.

### Evaluation of scLM using experimental data

To further demonstrate the performance of scLM, we compared scLM with the other methods (LTMG, CSN, Seurat-wgnca, MNN-wgcna, and SCENIC) on experimental scRNA-seq datasets. For comparisons, we used two in-house datasets from lung tumor and adjacent normal tissues as well as three public datasets from breast cancer (BR), head and neck squamous cell carcinoma (HNSCC), and melanoma. The data pre-processing procedures are described in the Methods section.

To assess and quantify clustering accuracy on real datasets, we used performance metrics including the Calinski-Harabasz (CH) index [38], Dunn index [40], and Davies-Bouldin (DB) index [39], to rank these methods. Importantly, scLM produced sets of clusters that showed significantly higher CH value than other methods (Figure 3A), especially higher than LTMG (P-value = 1.75E-07) and MNN-wgcna (P-value = 0.02), demonstrating that scLM achieves better cluster validity than other methods based on average between- and within-cluster sum of squares. In addition, compared to other methods, scLM also achieved significantly higher Dunn index scores representing better inter-cluster separation and intra-cluster compactness (Figure 3B), and lower DB index scores reflecting higher cluster quality (Figure 3C). Though SCENIC and Seurat-wgcna showed higher DB index score in the HNSCC dataset, they failed to show superior performance on the other datasets. Thus scLM proved to achieve the best partitioning of co-expressed gene clusters that are most distinct from each other.

**Figure 3.**
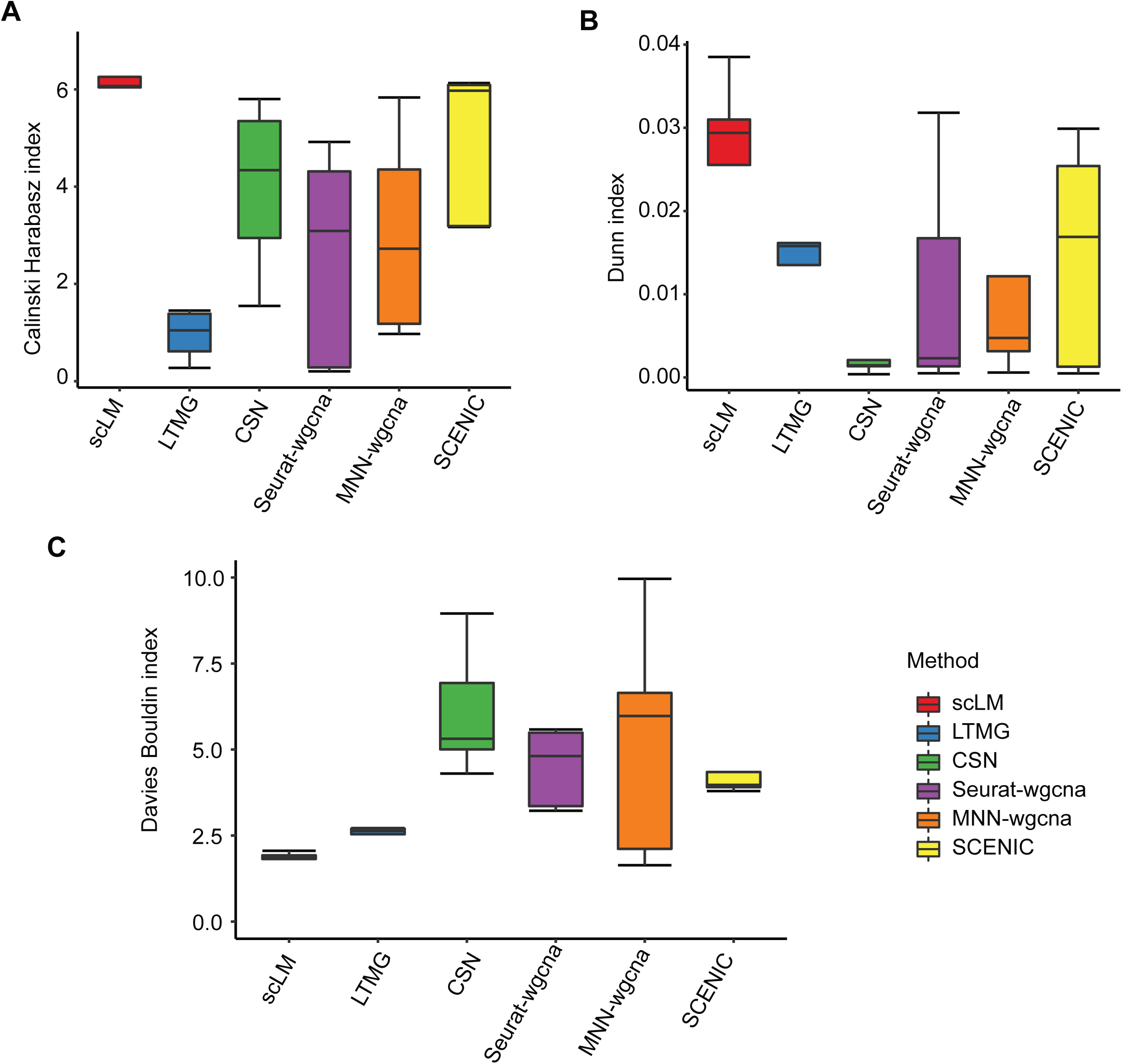
Evaluation of scLM using experimental data. scLM’s performance is compared against the other methods (LTMG, CSN, Seurat-wgcna, MNN-wgcna, and SCENIC) on five experimental datasets (BR, HNSCC, lungTumor, lungNormal, and Melanoma). Multiple evaluation indexes were used, including: **(A)** the Calinski-Harabasz (CH) index, **(B)** the Davies–Bouldin (DB) index, and **(C)** the Dunn index.

### scLM identified co-expressed genes with significantly enriched biological functions

As co-expressed genes are likely to be enriched with biological functions, we compared the extent to which different methods affect the functional discovery, based on their identified co-expressed genes. First, the aforementioned methods were evaluated for their capability to detect enriched GO terms in the five experimental datasets. Different methods identified gene clusters enriched with different GO enrichment results. The average number of significantly enriched GO terms (adjusted p-value < 0.05) ranges from 15 to 184 (Figure 4A). scLM extracted co-expressed genes with more enriched functional terms than other methods in each of the five datasets. SCENIC identified relatively high number of GO terms in BR dataset, whereas low number of enriched GO terms in the other datasets. In contrast, CSN identified relatively high number of GO terms in lungTumor dataset but low number of enriched GO terms in other datasets. LTMG, Seurat, and MNN all showed lower number of enriched GO terms in five datasets. Similar results were observed when we strengthened the enrichment significance by the adjusted p-value < 0.01 (**Figure S1**). The number of significant terms became fewer for all the methods, yet scLM identified the most on all the datasets. Some methods, like LTMG, failed to identify gene clusters with enriched terms at the threshold of adjusted p-value < 0.01.

**Figure 4.**
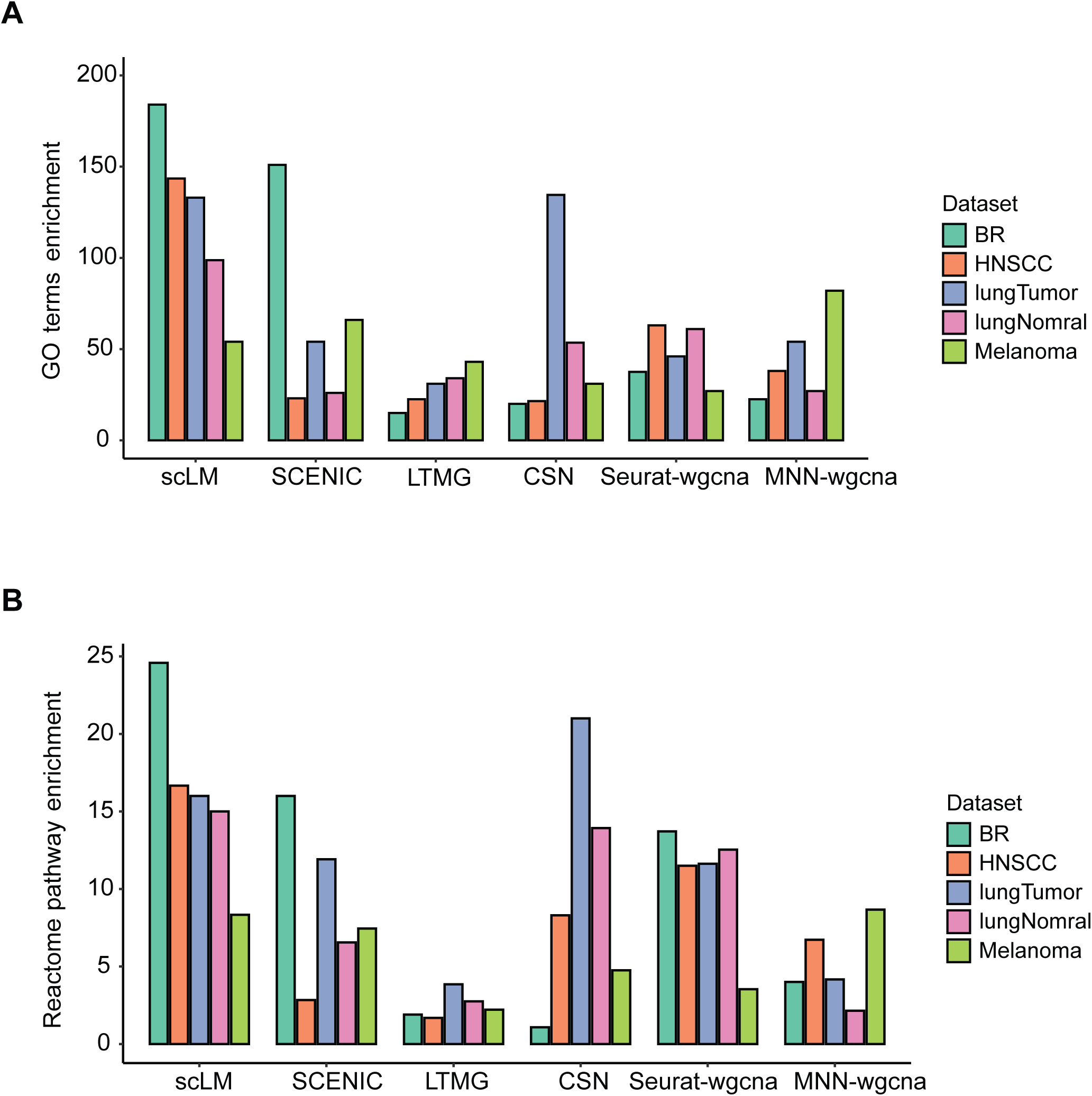
scLM identified co-expressed genes with significantly enriched biological functions. **A.** The average number of significantly enriched GO terms (adjusted *p*-value <= 0.05), based on the co-expressed genes identified by different methods. **B.** The average number of significantly enriched pathways (adjusted *p*-value <= 0.05), based on the co-expressed genes identified by different methods.

In addition to GO terms, we also examined the enriched pathways in the Reactome database, based on the co-expressed genes identified by different methods (Figure 4B). Different methods showed different pathway enrichment results. Importantly, scLM identified co-expressed genes with more enriched pathways than other methods in each of the five datasets. Though CSN identified relatively high number of pathways in lungTumor dataset, scLM shows prevalently more enriched pathways in all five datasets. Taken together, these results demonstrate that scLM outperforms other methods in functional discovery of co-expressed genes.

### scLM identified the tumor-specific modules enriched in specific cell state

In real-world scenarios, single cells from different patients or different data sources often demonstrate highly different cell numbers, largely due to strong batch effects and technical issues. The scLM method is designed to address such highly unbalanced data that could outperform other competitors on such datasets. To validate the effectiveness of scLM, we intentionally selected patient samples that varied with respect to cell number, which could create challenges for this method. As a case study, we used scLM to analyze our in-house scRNA-seq profiling from 4 NSCLC patients (P1-P4) [35] to identify the co-expressed genes in tumor and normal epithelial cells respectively. In tumor cells (Figure 5A, heatmap of latent variables), we discovered 12 co-expressed gene modules in the latent space (T1 – T12). These modules showed clear differences but were consistently concordant across patients (Figure 5A, heatmaps P1 – P4), even though the single cells from different patients presented strong heterogeneity and batch effects (Figure 5B, left panel).

**Figure 5.**
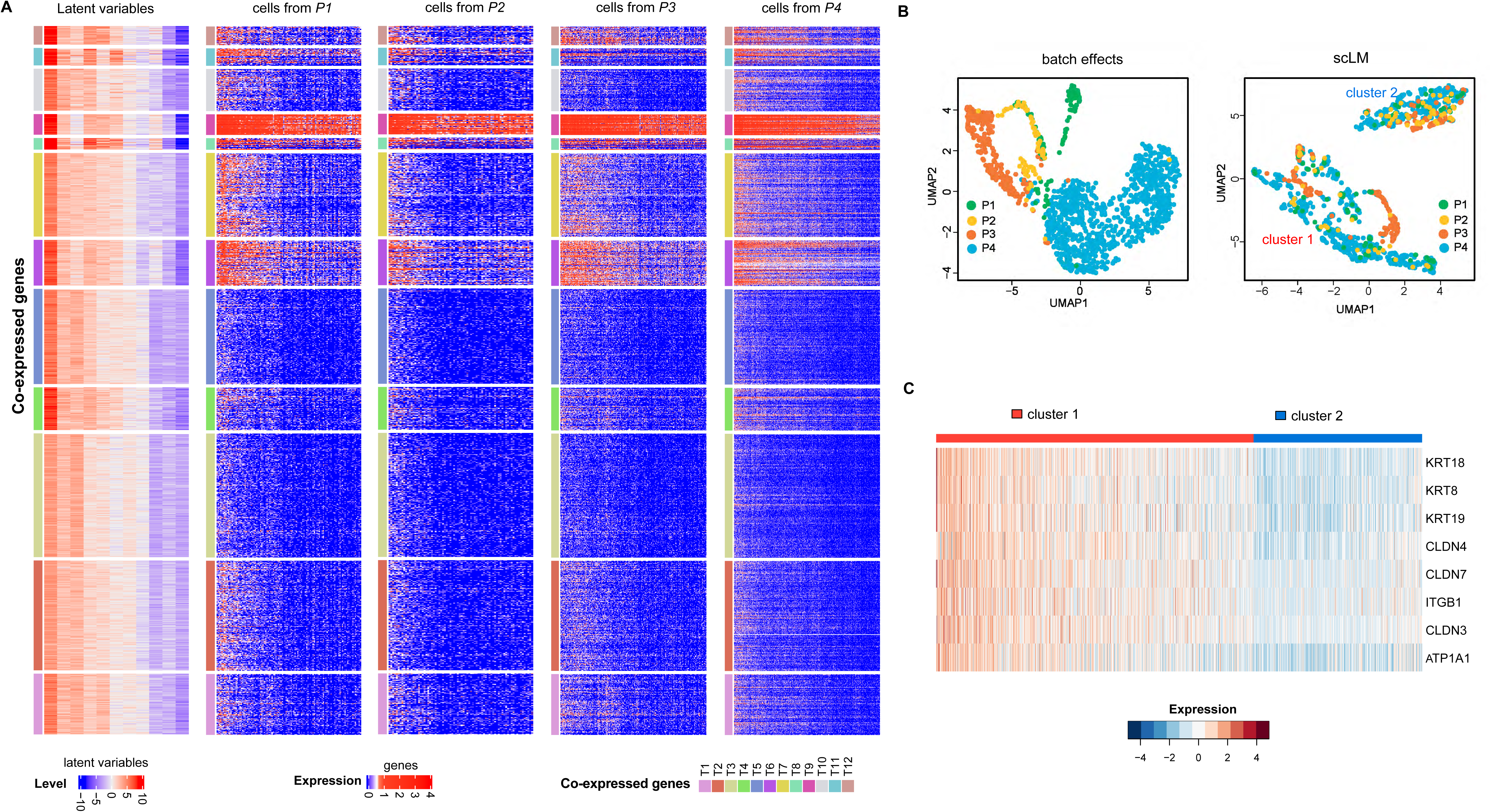
scLM identified co-expression modules that characterize subtle cell subpopulations. **A.** Simultaneous and consensus clustering of genes across lung tumor cells from four patients. ScLM reveals 12 co-expressed gene modules characterized by the latent variables as well as shown in real gene expression data across patients. In each heatmap, rows are genes assigned to 12 co-expression modules. In each co-expression module, genes are consistently over-expressed (red) or under-expressed (blue). **B.** The left panel shows strong batch effects of different patients. The right panel depicts the UMAP visualization of single cells characterized by the co-expression modules of scLM. Two evenly distributed clusters (cluster 1 and cluster 2) are identified. Different patients are distinguished by colors. **C.** Heatmap showing the differential expression pattern of EMT-related genes in two clusters.

Using the 12 co-expression modules, the single cells were separated into two major clusters. In each cluster, cells from different patients mixed well without interference from batch effects (Figure 5B, right panel), which further support that the co-expression modules are consistent across patients. Interestingly, we found that cluster 1 had higher expression of epithelial functional markers (EMT-related genes) than cluster 2 (Figure 5C). These results indicate that co-expression modules are capable of characterizing specific cell phenotypes.

Similarly, in normal single cells, we observed 13 co-expressed gene modules (N1 – N13) that showed concordant expression across individual patients (**Figure S2**). Then we compared the co-expression modules identified from tumor and normal cells. Four modules (T1, T3, T4, and T10) were not correlated with any normal modules, suggesting that they were tumor-specific (**Supplementary file, Figure S3**).

### scLM identified a common program across three types of cancer

To explore the common mechanism of different cancers, we next extended the application of scLM to HNSCC and melanoma. In addition to the 12 co-expression modules in NSCLC, we identified 11 modules in HNSCC and 14 modules in melanoma. To determine the similarities of these co-expression modules, we performed a pair-wise comparison using weighted jaccard similarity, followed by hierarchical clustering. As shown in the diagram (**Figure S4A**), we found that most branches were dominated by a mixture of cancer types. Importantly, we identified a branch with high similarity among lung_m9, hnscc_m7, and melanoma_m12 modules.

These three similar modules substantially overlapped with 91 genes, which were defined as a common program across three cancer types. To gain insights into the biological functions of the common program, we performed enrichment analysis in the Hallmark database (Figure S4B). The MYC targets v1 and hypoxia were the top enriched terms, involving the genes *FOS*, *GAPDH*, *HLA-A*, and *NFKB1A*. This result suggests the common oncogenesis mechanism regardless of cancer types (**Supplementary file**).

From the applications of scLM, we see three usefulness of identifying co-expressed genes from multiple datasets: (1) Co-expressed genes can reveal novel biological processes. An example is the lung tumor specific module that highlights cell-cell communication in tumor micro-environment (Supplementary file, Figure S3C). (2) Co-expressed genes contribute to the subtle characterization of single cell states. (3) With the co-expressed genes identified, both specific and common gene modules can further be explored. Examples are the lung tumor specific module and the common co-expression module of three different cancer types (Supplementary file, Figure S4A).

Given the merits of scLM performance, several potential limitations warrant further study. First, zero-inflated genes are excluded during pre-processing. The main reason is that, genes with inflated zeros are not informative and have negligible meaningful contribution to co-expression. The other reason is, with the fast advance of scRNA-seq technology, zero-inflation issue will be very minimal in near future. Second, in future work, we will examine the necessity of providing zero-inflation models, which specifically deal with data of poor sequencing depth and strong dropout effects. Third, the computational cost of our method can be further reduced. We have already utilized Rcpp programming and parallel computing to dramatically boost the performance of scLM. However, considering scRNA-seq data is growing into million-cell level, we will explore the use of GPU computing and cloud-based approaches to catch up the scale of the future scRNA-seq data.

## Conclusion

Genes with coordinate biological functions are frequently co-transcribed resulting in correlated expression profiles. scRNA-seq technologies enable the systematic interrogation of gene co-expression modules within and across distinct cell types, and within and across distinct samples. In this study, we introduce a novel method (scLM) to identify co-expressed genes across multiple single-cell datasets simultaneously. To our knowledge, scLM is the first available tool that is capable of leveraging multiple scRNA-seq datasets to accurately detect co-expressed genes. scLM uses the conditional negative binomial distribution with latent variables to disentangle co-expression patterns across multiple datasets.

Compared with other methods, scLM has several key advantages: (1) scLM accounts for data heterogeneity and variances among multiple datasets, such as unbalanced sequencing depths and technical biases in library preparation. (2) scLM leverages information across datasets for detecting stable and conserved modules with high accuracy and reproducibility. (3) scLM is an integrated pipeline that uses only raw count matrix without prior batch-correction and thus, can be easily and rapidly applied the scRNA-seq data output. Overall, scLM opens possibilities for further investigation and mechanistic interpretation of co-expressed gene modules. With the growing applications of scRNA-seq, scLM is poised to become a valuable tool for elucidating co-expression patterns in single cell transcriptomic studies.

## Ethical statements

For our in-house data, fresh remnant tumor and adjacent normal tissues were collected at the time of elective curative resection by the Tumor Tissue and Pathology Shared Resource (TTPSR) of the Wake Forest Baptist Medical Center Comprehensive Cancer Center (WFBMC-CCC). Collections by the TTPSR adhere to Institutional Review Board approved procedures (Advanced Tumor Bank protocol CCCWFU 01403, TTPSR collections IRB BG04-104 which also allows for the use of de-identified protected health information along with the tissue samples).

Acquisition of de-identified samples from the TTPSR for single cell isolation and research use was in accordance with approved IRB protocol 00048977.

## Authors’ contributions

WZ, JS and QS developed the structure and arguments for the paper. JS and QS wrote the manuscript, with input from LM and WZ. All the authors reviewed and approved the final manuscript.

## Acknowledgements

The authors thank Elizabeth Forbes and Karen Klein for professional editing service. This work was supported in part by the Cancer Genomics, Tumor Tissue Repository, and Bioinformatics Shared Resources under the NCI Cancer Center Support Grant to the Comprehensive Cancer Center of Wake Forest University Health Sciences (P30CA012197). WZ is supported by the Hanes and Willis Professorship in Cancer. Additional support for QS and WZ were provided by a Fellowship to WZ from the National Foundation for Cancer Research.

## Supplementary material

**Figure S1. Significant GO terms enriched in co-expressed genes by different methods**

The average number of GO terms detected as significantly enriched with tighter threshold (adjusted p.value < 0.01), based on the co-expressed genes identified by different methods.

**Figure S2. scLM identified co-expression modules in lung normal cells**

scLM yielded 13 co-expressed gene modules characterized by the latent variables as well as in gene expression data. In each heatmap, rows are genes assigned to 13 co-expression modules. In each co-expression module, genes are consistently over-expressed (red) or under-expressed (blue).

**Figure S3. scLM uncovered tumor-specific modules enriched in specific cell subpopulations**

**A.** Heatmap depicts the pairwise correlations between the 12 co-expression modules from tumor (rows) and the 13 co-expression modules from normal (columns). Red color with star represents significant association with Pearson correlation > 0.5. **B.** UMAP visualization of single cells labeled by high versus low expression levels of the tumor-specific module T10. **C.** Significantly enriched pathways in the REACTOME database are identified based on the tumor-specific modules (T1, T3, T10). The x-axis represents the enrichment score defined by –log10(adjusted p-value) of the enrichment test. **D.** Putative upstream regulators of T10 were identified and labeled as blue. E. Kaplan-Meier survival curves for patients with lung squamous cell cancer (LUSC, n = 545), stratified by average expression (top 30% versus bottom 30%) of the tumor-specific module T10. Log-rank test p-value is shown. The y-axis represents the probability of overall survival, and the x axis is time in Months.

**Figure S4. scLM identified a common program across three cancer types**

**A.** The dendrogram presents the similarity of the co-expression modules from three cancer types, including 12 co-expression modules from NSCLC, 11 from HNSCC, and 14 from melanoma. **B.** Bar plots show the significantly enriched processes of the common program in Hallmark database. The x-axis represents the enrichment score defined by *–log10(adj.P-value)* of the enrichment test.

**Figure S5. Tumor-specific modules enriched in specific cell subpopulations**

UMAP visualization of single cells labeled by different expression level (high v.s. low) of the tumor-specific modules (T1, T3, T4).

**Figure S6. Biological mechanism underlying the tumor-specific modules**

A-B. Significantly enriched processes in the KEGG (**A**) and Hallmark (**B**) databases are identified based on the tumor-specific modules (T1, T3, T4, and T10). The x-axis represents the enrichment score defined by *–log10(adj.P-value)* of the enrichment test. **C.** Putative upstream regulators of the tumor-specific modules (T1, T3, and T4) were identified and labeled as blue.

**Figure S7. Prognostic significance of the tumor-specific modules**

**A.** Kaplan-Meier survival curves for LUSC patients, which are stratified by the average expression (top 30% versus bottom 30%) of tumor-specific modules (T1, T3, and T4). **B.** Kaplan-Meier survival curves for LUAD patients, which are stratified by the average expression (top 30% versus bottom 30%) of tumor-specific modules (T1, T3, T4, and T10). Log-rank test p-value is shown. The y-axis represents the probability of overall survival, and the x axis is time in Months/Days.

**Figure S8. A common program across three cancer types**

**A.** The left panel shows the strong batch effects of different patients. The right panel depicts the UMAP visualization of single cells characterized by co-expression modules. Different patients are distinguished by color. The upper part shows the melanoma data, whereas the lower part shows the HNSCC data. **B.** The dendrogram presents the similarity of the co-expression modules from three cancer types, including 12 co-expression modules from NSCLC, 11 from HNSCC, and 14 from melanoma. **C.** Boxplots shows the expression levels of the three modules (hnscc_m7, lung_m9, melanoma_m12) in their corresponding tumor samples versus normal samples from TCGA. Kaplan-Meier (KM) curves show that hnscc_m7 and melanoma_m12 were associated with poor overall survival in HNSCC and melanoma patients, respectively. **D.** Bar plots show the significantly enriched processes of the common program in Hallmark database. The x-axis represents the enrichment score defined by -log10(adj.p-value) of the enrichment test. **Figure S9. Co-expression modules are potential markers for immunothreapy response** Melanoma modules 5 and 9 distinguished partial response (PR) patients from the complete response (CR) patients in an external cohort that received immunotherapy treatment.

## References

[1] Ferrari R, Forabosco P, Vandrovcova J, Warren JD, Momeni P, Weale ME, et al. Frontotemporal Dementia: insights into the biological underpinnings of disease through gene co-expression network analysis. Neuropathology and Applied Neurobiology 2016;42:38-.

[2] Yang Y, Han L, Yuan Y, Li J, Hei N, Liang H. Gene co-expression network analysis reveals common system-level properties of prognostic genes across cancer types. Nat Commun 2014;5:3231.

[3] Stuart JM, Segal E, Koller D, Kim SK. A gene-coexpression network for global discovery of conserved genetic modules. Science 2003;302:249–55.

[4] Jerby-Arnon L, Shah P, Cuoco MS, Rodman C, Su MJ, Melms JC, et al. A Cancer Cell Program Promotes T Cell Exclusion and Resistance to Checkpoint Blockade. Cell 2018;175:984–97 e24.

[5] Singer M, Wang C, Cong L, Marjanovic ND, Kowalczyk MS, Zhang H, et al. A Distinct Gene Module for Dysfunction Uncoupled from Activation in Tumor-Infiltrating T Cells. Cell 2017;171:1221–3.

[6] Puram SV, Tirosh I, Parikh AS, Patel AP, Yizhak K, Gillespie S, et al. Single-Cell Transcriptomic Analysis of Primary and Metastatic Tumor Ecosystems in Head and Neck Cancer. Cell 2017;171:1611–24 e24.

[7] Chihara N, Madi A, Kondo T, Zhang H, Acharya N, Singer M, et al. Induction and transcriptional regulation of the co-inhibitory gene module in T cells. Nature 2018;558:454–9.

[8] Lawson DA, Bhakta NR, Kessenbrock K, Prummel KD, Yu Y, Takai K, et al. Single-cell analysis reveals a stem-cell program in human metastatic breast cancer cells. Nature 2015;526:131–5.

[9] Svensson V. Droplet scRNA-seq is not zero-inflated. Nature Biotechnology 2020;38:147–50.

[10] Lun AT, Bach K, Marioni JC. Pooling across cells to normalize single-cell RNA sequencing data with many zero counts. Genome Biol 2016;17:75.

[11] Vieth B, Ziegenhain C, Parekh S, Enard W, Hellmann I. powsimR: power analysis for bulk and single cell RNA-seq experiments. Bioinformatics 2017;33:3486–8.

[12] Grun D, Kester L, van Oudenaarden A. Validation of noise models for single-cell transcriptomics. Nat Methods 2014;11:637–40.

[13] Bacher R, Kendziorski C. Design and computational analysis of single-cell RNA-sequencing experiments. Genome Biol 2016;17:63.

[14] Marinov GK, Williams BA, McCue K, Schroth GP, Gertz J, Myers RM, et al. From single-cell to cell-pool transcriptomes: stochasticity in gene expression and RNA splicing. Genome Res 2014;24:496–510.

[15] Kolodziejczyk AA, Kim JK, Svensson V, Marioni JC, Teichmann SA. The technology and biology of single-cell RNA sequencing. Mol Cell 2015;58:610–20.

[16] Macaulay IC, Voet T. Single cell genomics: advances and future perspectives. PLoS Genet 2014;10:e1004126.

[17] Azizi E, Carr AJ, Plitas G, Cornish AE, Konopacki C, Prabhakaran S, et al. Single-Cell Map of Diverse Immune Phenotypes in the Breast Tumor Microenvironment. Cell 2018;174:1293–308 e36.

[18] Cusanovich DA, Hill AJ, Aghamirzaie D, Daza RM, Pliner HA, Berletch JB, et al. A Single-Cell Atlas of In Vivo Mammalian Chromatin Accessibility. Cell 2018;174:1309–24 e18.

[19] Muraro MJ, Dharmadhikari G, Grun D, Groen N, Dielen T, Jansen E, et al. A Single-Cell Transcriptome Atlas of the Human Pancreas. Cell Syst 2016;3:385–94 e3.

[20] Tabula Muris C, Overall c, Logistical c, Organ c, processing, Library p, et al. Single-cell transcriptomics of 20 mouse organs creates a Tabula Muris. Nature 2018;562:367–72.

[21] Buenrostro JD, Corces MR, Lareau CA, Wu B, Schep AN, Aryee MJ, et al. Integrated Single-Cell Analysis Maps the Continuous Regulatory Landscape of Human Hematopoietic Differentiation. Cell 2018;173:1535–48 e16.

[22] Langfelder P, Horvath S. WGCNA: an R package for weighted correlation network analysis. BMC Bioinformatics 2008;9:559.

[23] Abu-Jamous B, Kelly SJGb. Clust: automatic extraction of optimal co-expressed gene clusters from gene expression data2018;19:172.

[24] Aibar S, González-Blas CB, Moerman T, Huynh-Thu VA, Imrichova H, Hulselmans G, et al. SCENIC: single-cell regulatory network inference and clustering. Nature Methods 2017;14:1083–6.

[25] Dai H, Li L, Zeng T, Chen L. Cell-specific network constructed by single-cell RNA sequencing data. Nucleic Acids Res 2019;47:e62.

[26] Wan C, Chang W, Zhang Y, Shah F, Lu X, Zang Y, et al. LTMG: a novel statistical modeling of transcriptional expression states in single-cell RNA-Seq data. Nucleic Acids Res 2019;47:e111.

[27] Raj A, Peskin CS, Tranchina D, Vargas DY, Tyagi S. Stochastic mRNA synthesis in mammalian cells. PLoS biology 2006;4.

[28] Brennecke P, Anders S, Kim JK, Kolodziejczyk AA, Zhang X, Proserpio V, et al. Accounting for technical noise in single-cell RNA-seq experiments. Nat Methods 2013;10:1093–5.

[29] Anders S, Huber W. Differential expression analysis for sequence count data. Genome Biol 2010;11:R106.

[30] McCarthy DJ, Chen Y, Smyth GK. Differential expression analysis of multifactor RNA-Seq experiments with respect to biological variation. Nucleic Acids Res 2012;40:4288–97.

[31] Love MI, Huber W, Anders S. Moderated estimation of fold change and dispersion for RNA-seq data with DESeq2. Genome Biol 2014;15:550.

[32] Hafemeister C, Satija R. Normalization and variance stabilization of single-cell RNA-seq data using regularized negative binomial regression. Genome Biology 2019;20.

[33] Wang Z, Ma S, Zappitelli M, Parikh C, Wang CY, Devarajan P. Penalized count data regression with application to hospital stay after pediatric cardiac surgery. Stat Methods Med Res 2016;25:2685–703.

[34] Zappia L, Phipson B, Oshlack A. Splatter: simulation of single-cell RNA sequencing data. Genome Biol 2017;18:174.

[35] Song Q, Hawkins GA, Wudel L, Chou PC, Forbes E, Pullikuth AK, et al. Dissecting intratumoral myeloid cell plasticity by single cell RNA-seq. Cancer Med 2019.

[36] Tirosh I, Izar B, Prakadan SM, Wadsworth MH, 2nd, Treacy D, Trombetta JJ, et al. Dissecting the multicellular ecosystem of metastatic melanoma by single-cell RNA-seq. Science 2016;352:189–96.

[37] Hubert L, Arabie PJJoc. Comparing partitions1985;2:193–218.

[38] Caliński T, Harabasz JJCiS-t, Methods. A dendrite method for cluster analysis1974;3:1–27.

[39] Davies DL, Bouldin DWJItopa, intelligence m. A cluster separation measure1979:224–7.

[40] Dunn JC. Well-separated clusters and optimal fuzzy partitions. Journal of cybernetics 1974;4:95–104.

[41] Krijthe JJCS. Rtsne: T-distributed stochastic neighbor embedding using a Barnes-Hut implementation. R package version 0.132015.

[42] Liberzon A, Subramanian A, Pinchback R, Thorvaldsdottir H, Tamayo P, Mesirov JP. Molecular signatures database (MSigDB) 3.0. Bioinformatics 2011;27:1739–40.

[43] Fabregat A, Jupe S, Matthews L, Sidiropoulos K, Gillespie M, Garapati P, et al. The Reactome Pathway Knowledgebase. Nucleic Acids Res 2018;46:D649–D55.

[44] Du J, Yuan Z, Ma Z, Song J, Xie X, Chen Y. KEGG-PATH: Kyoto encyclopedia of genes and genomes-based pathway analysis using a path analysis model. Mol Biosyst 2014;10:2441–7.

[45] Yu GC, Wang LG, Han YY, He QY. clusterProfiler: an R Package for Comparing Biological Themes Among Gene Clusters. Omics-a Journal of Integrative Biology 2012;16:284–7.

[46] Stuart T, Butler A, Hoffman P, Hafemeister C, Papalexi E, Mauck WM, 3rd, et al. Comprehensive Integration of Single-Cell Data. Cell 2019;177:1888–902 e21.

[47] Haghverdi L, Lun ATL, Morgan MD, Marioni JC. Batch effects in single-cell RNA-sequencing data are corrected by matching mutual nearest neighbors. Nat Biotechnol 2018;36:421–7.

