## Supplementary Figures for "scLM: automatic detection of consensus gene clusters across multiple single-cell datasets"

**Figure S1**

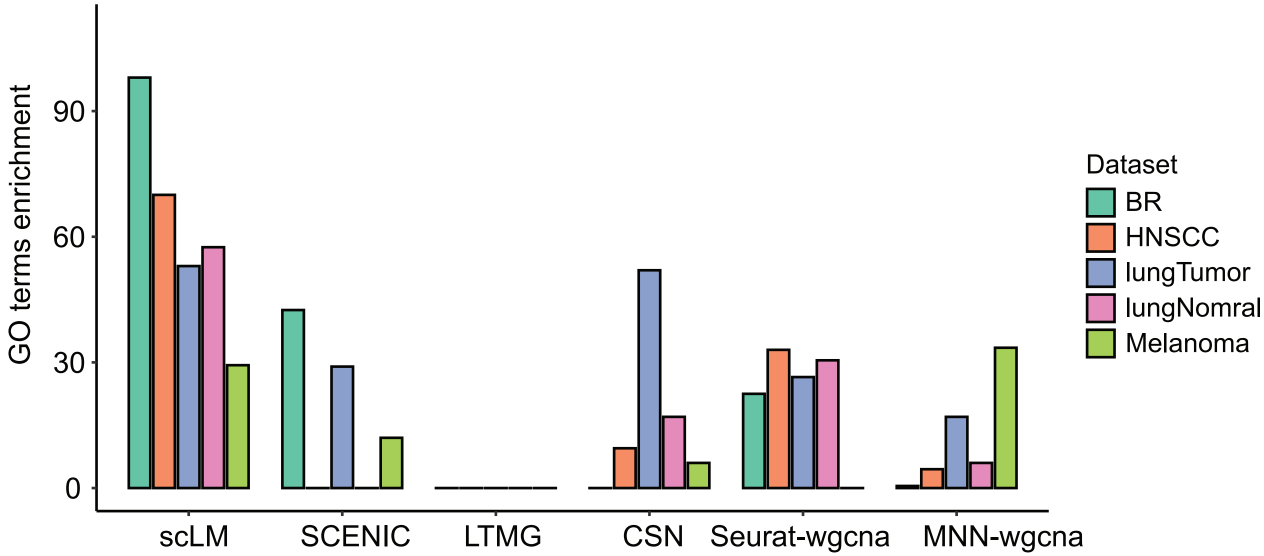
GO terms enrichment
90
60
30
0
scLM
SCENIC
LTMG
CSN
Seurat-wgcna
MNN-wgcna
Dataset
BR
HNSCC
lungTumor
lungNomral
Melanoma

#### Figure S2

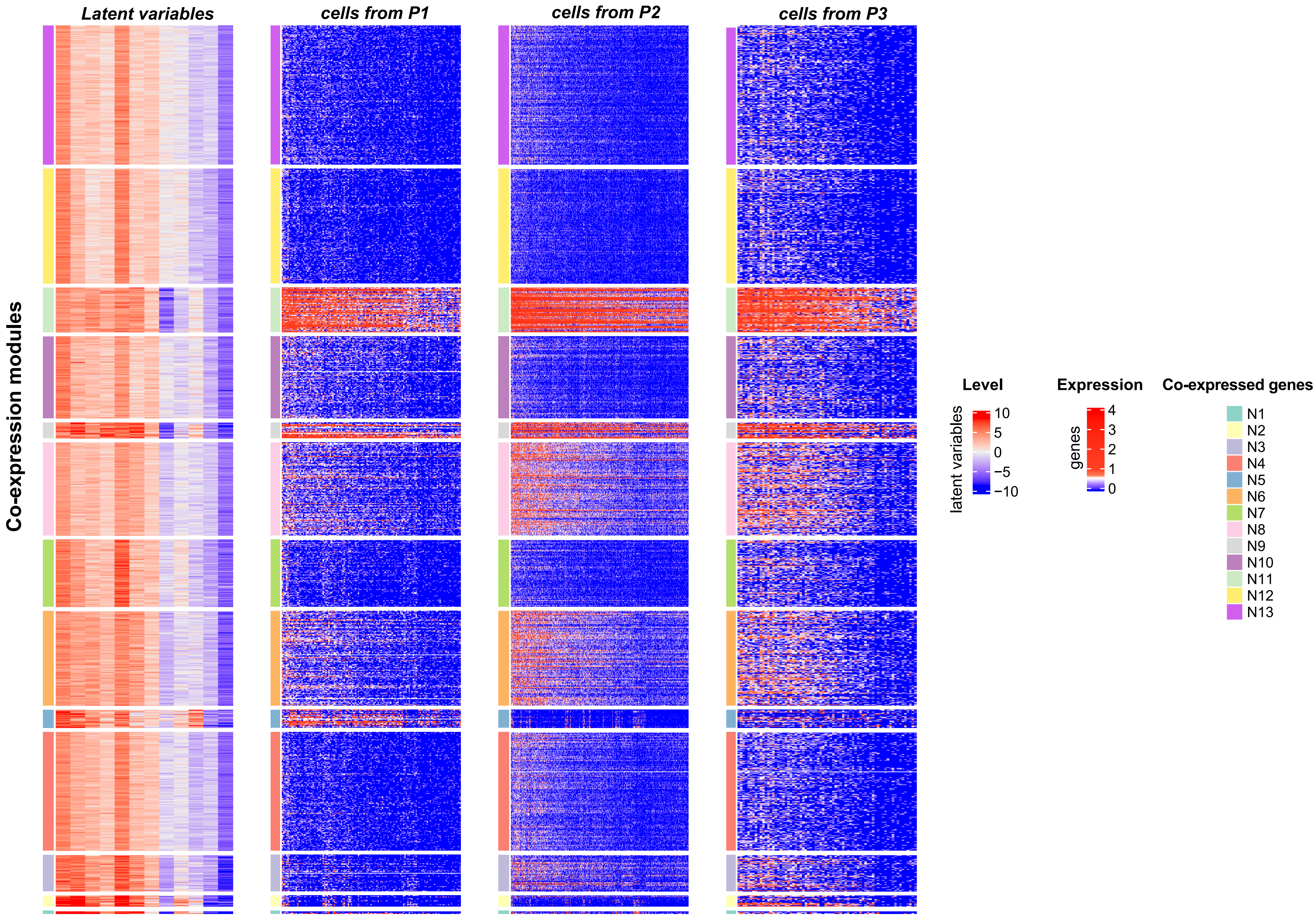


**Figure S2. scLM identified co-expression modules in lung normal cells**

Simultaneous clustering of normal cells across three patients. ScLM yielded 13 co-expression gene modules characterized by the latent variables as well as in real gene expression data across patients. In each heatmap, rows are genes assigned to 13 co-expression modules. In each co-expression module, genes are consistently over-expressed (red) or under-expressed (blue).

Figure S3

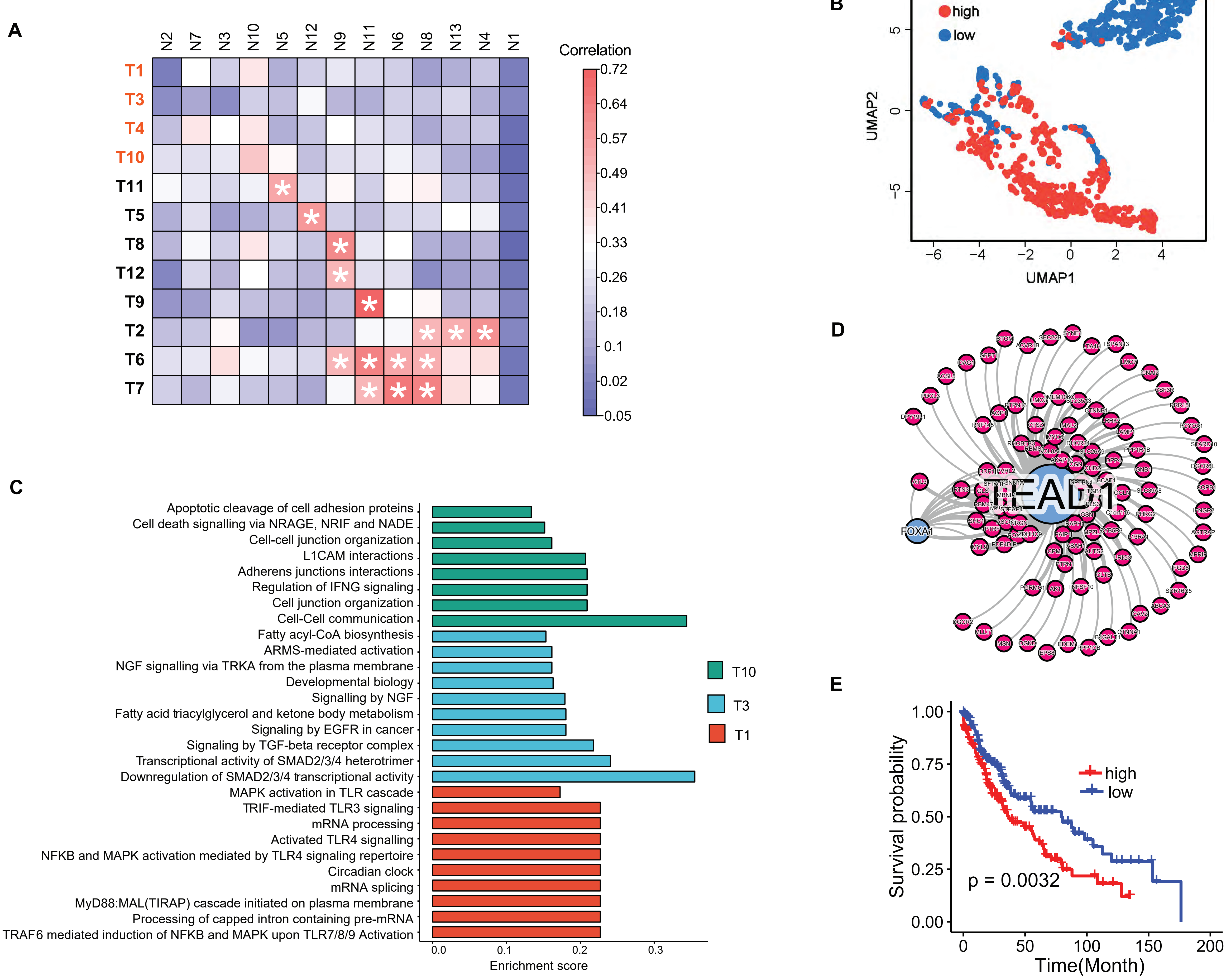


**Figure S3. scLM uncovered tumor-specific modules enriched in specific cell subpopulations**

**A.** Heatmap depicts the pairwise correlations between the 12 co-expression modules from tumor (rows) and the 13 co-expression modules from normal (columns). Red color with star represents significant association with Pearson correlation >0.5.

**B.** UMAP visualization of single cells labeled by high versus low expression levels of the tumor-specific module T10.

**C.** Significantly enriched pathways in the REACTOME database are identified based on the tumor-specific modules (T1, T3, T10). The x-axis represents the enrichment score defined by –log10(adjusted p- value) of the enrichment test.

**D.** Putative upstream regulators of T10 were identified and labeled as blue. E. Kaplan-Meier survival curves for patients with lung squamous cell cancer (LUSC, n = 545), stratified by average expression (top 30% versus bottom 30%) of the tumor-specific module T10. Log-rank test p-value is shown. The y-axis represents the probability of overall survival, and the x axis is time in Months.

**Figure S4**

**A**

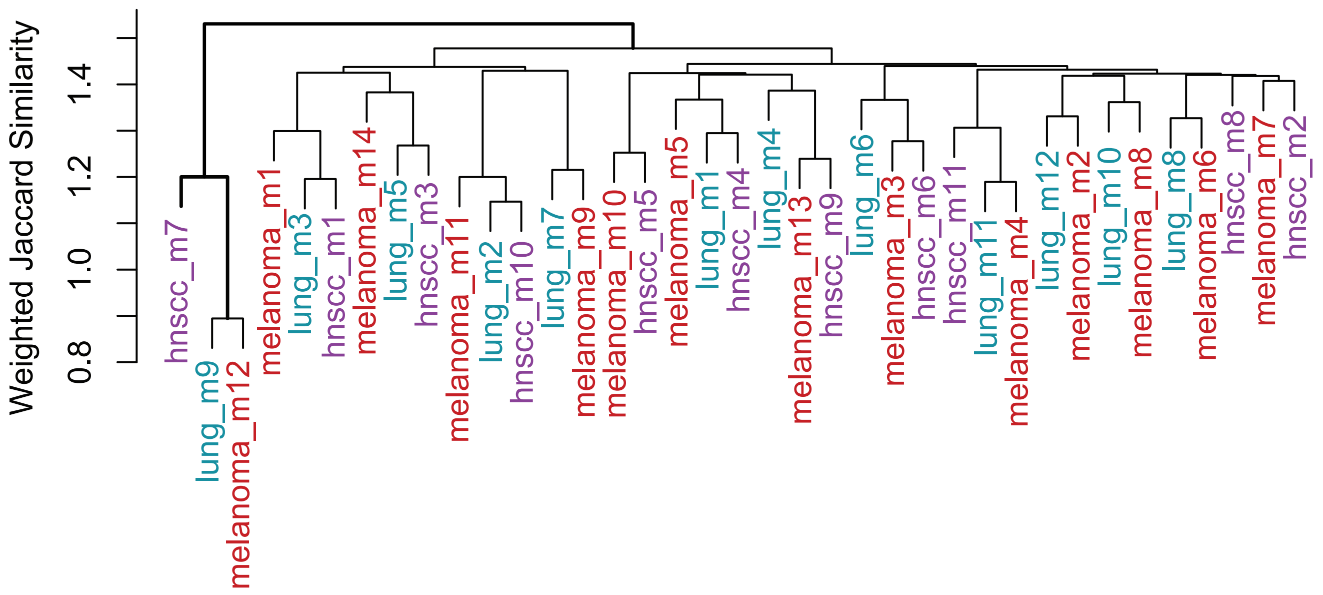
Weighted Jaccard Similarity
0.8
1.0
1.2
1.4
hnscc_m7
lung_m9
melanoma_m12
melanoma_m1
lung_m3
hnscc_m1
melanoma_m14
lung_m5
hnscc_m3
melanoma_m11
lung_m2
hnscc_m10
lung_m7
melanoma_m9
melanoma_m10
hnscc_m5
melanoma_m5
lung_m1
hnscc_m4
lung_m4
melanoma_m13
hnscc_m9
lung_m6
melanoma_m3
hnscc_m6
hnscc_m11
lung_m11
melanoma_m4
lung_m12
melanoma_m2
lung_m10
melanoma_m8
lung_m8
melanoma_m6
hnscc_m8
melanoma_m7
hnscc_m2

**B**

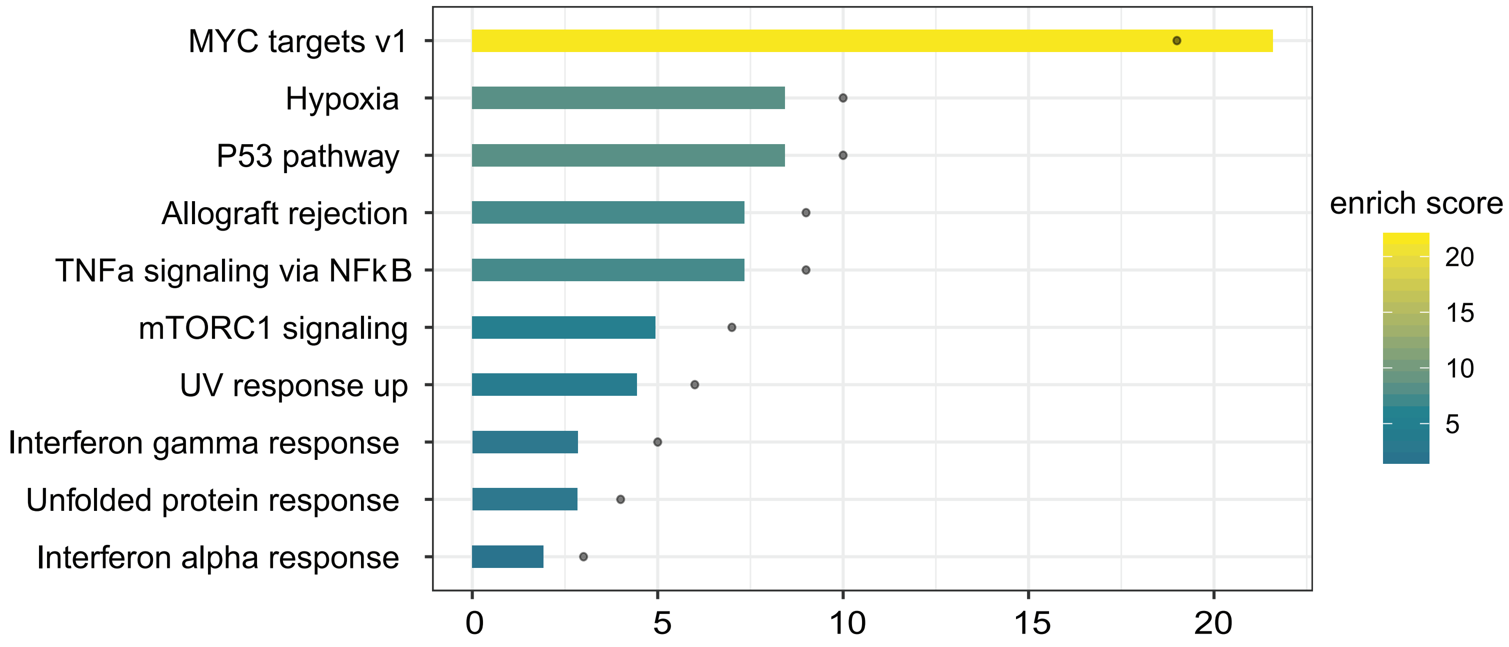
MYC targets v1
Hypoxia
P53 pathway
Allograft rejection
TNFa signaling via NFkB
mTORC1 signaling
UV response up
Interferon gamma response
Unfolded protein response
Interferon alpha response
0
5
10
15
20
enrich score
20
15
10
5

### Figure S5

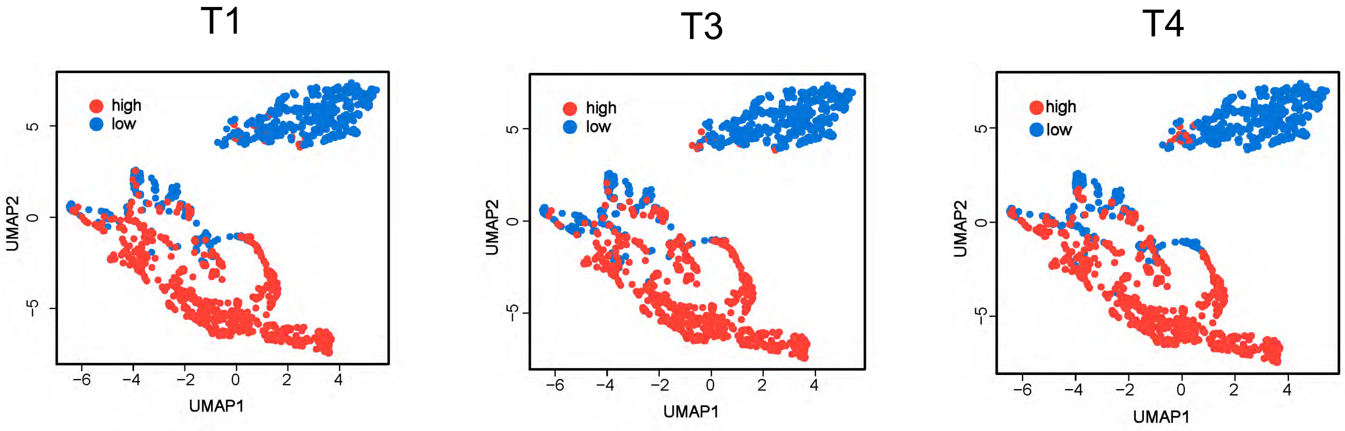
T1
high
low
UMAP2
UMAP1
5
0
−5
−6
−4
−2
0
2
4
T3
high
low
UMAP2
UMAP1
5
0
−5
−6
−4
−2
0
2
4
T4
high
low
UMAP2
UMAP1
5
0
−5
−6
−4
−2
0
2
4

**Figure S6**

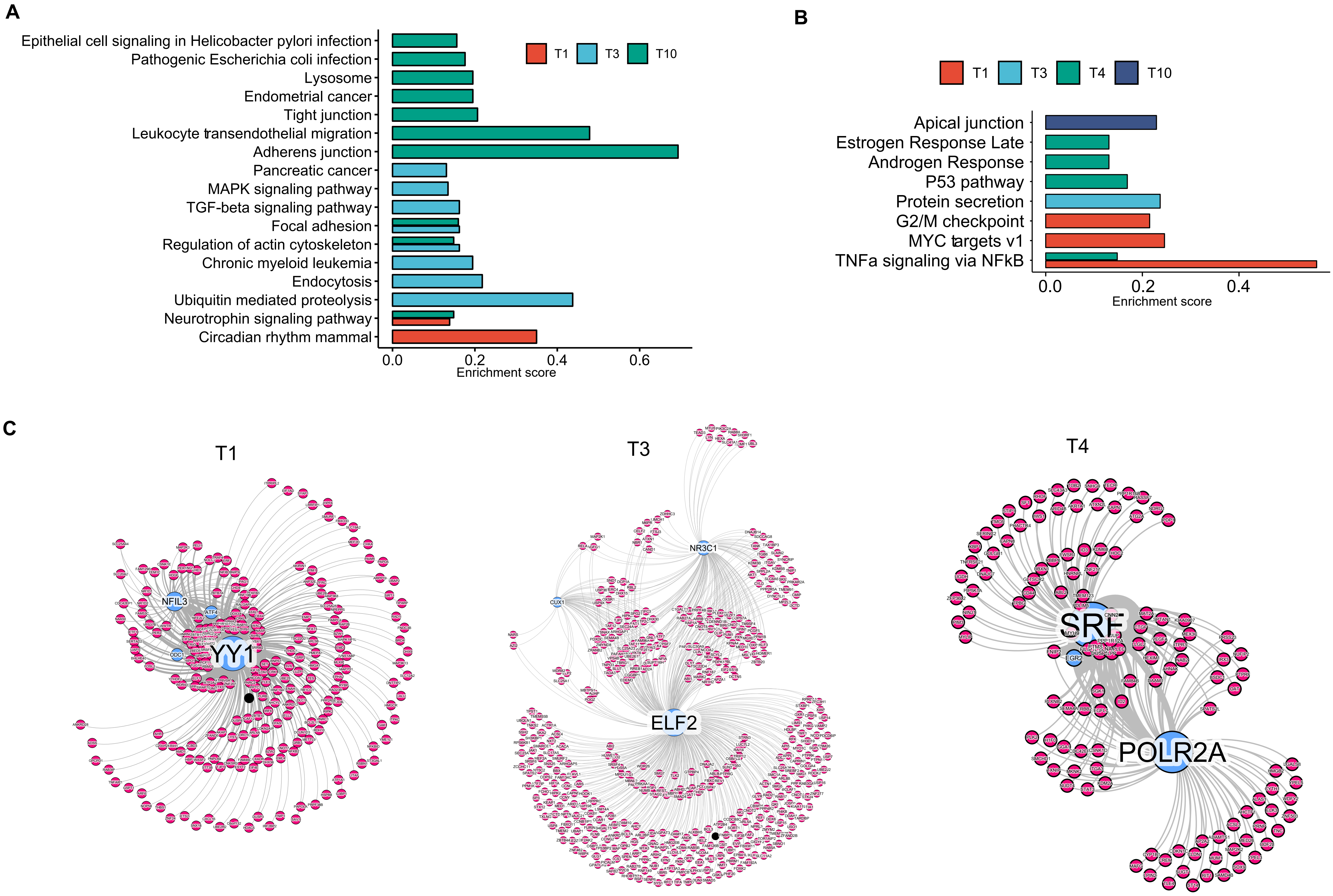
A
Epithelial cell signaling in Helicobacter pylori infection
Pathogenic Escherichia coli infection
Lysosome
Endometrial cancer
Tight junction
Leukocyte transendothelial migration
Adherens junction
Pancreatic cancer
MAPK signaling pathway
TGF-beta signaling pathway
Focal adhesion
Regulation of actin cytoskeleton
Chronic myeloid leukemia
Endocytosis
Ubiquitin mediated proteolysis
Neurotrophin signaling pathway
Circadian rhythm mammal
T1
T3
T10
0.0
0.2
0.4
0.6
Enrichment score
B
T1
T3
T4
T10
Apical junction
Estrogen Response Late
Androgen Response
P53 pathway
Protein secretion
G2/M checkpoint
MYC targets v1
TNFa signaling via NFkB
0.0
0.2
0.4
Enrichment score
C
T1
YY1
NFIL3
ATF4
ODC1
T3
NR3C1
CUX1
ELF2
T4
SRF
EGR2
POLR2A

**Figure S7**

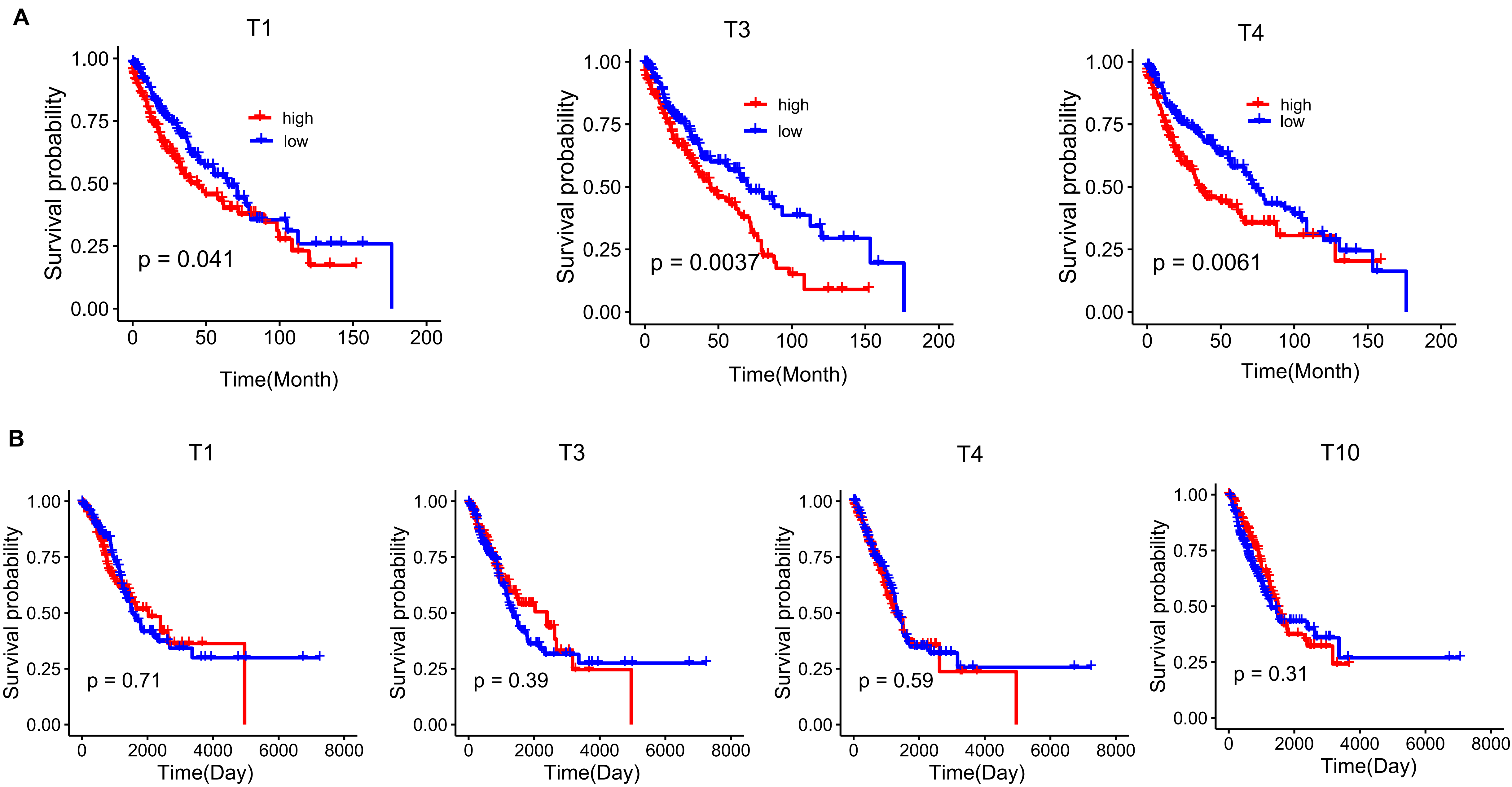
A
T1
Survival probability
1.00
0.75
0.50
0.25
0.00
high
low
p = 0.041
0
50
100
150
200
Time(Month)
T3
Survival probability
high
low
p = 0.0037
Time(Month)
T4
Survival probability
high
low
p = 0.0061
Time(Month)
B
T1
Survival probability
p = 0.71
0
2000
4000
6000
8000
Time(Day)
T3
Survival probability
p = 0.39
Time(Day)
T4
Survival probability
p = 0.59
Time(Day)
T10
Survival probability
p = 0.31
Time(Day)

**Figure S8**

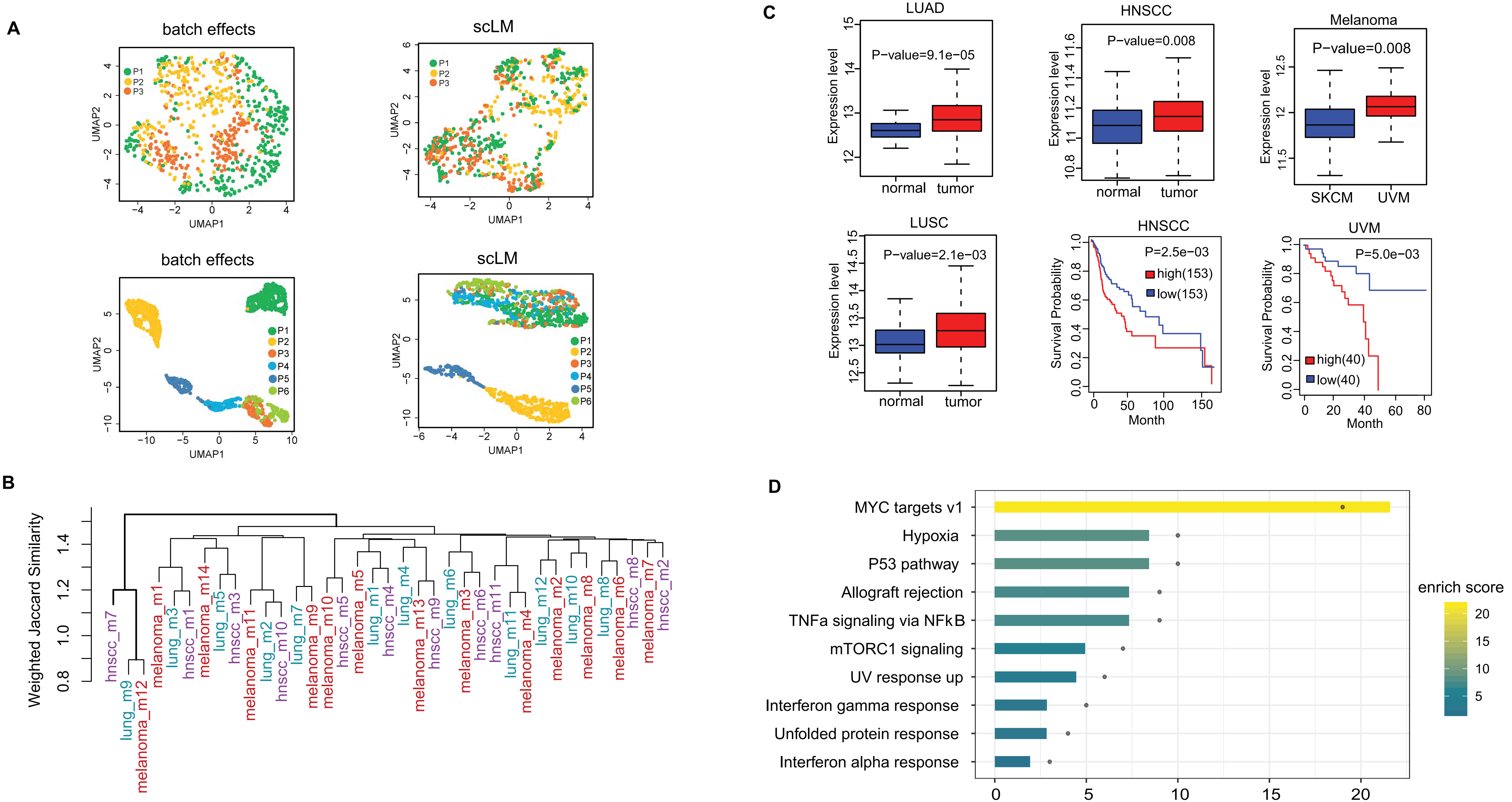
A
batch effects
scLM
batch effects
scLM
UMAP1
UMAP2
P1
P2
P3
P4
P5
P6
B
Weighted Jaccard Similarity
C
LUAD
P-value=9.1e−05
HNSCC
P-value=0.008
Melanoma
P-value=0.008
LUSC
P-value=2.1e−03
Expression level
normal
tumor
SKCM
UVM
HNSCC
P=2.5e−03
high(153)
low(153)
UVM
P=5.0e−03
high(40)
low(40)
Survival Probability
Month
D
MYC targets v1
Hypoxia
P53 pathway
Allograft rejection
TNFa signaling via NFkB
mTORC1 signaling
UV response up
Interferon gamma response
Unfolded protein response
Interferon alpha response
enrich score

#### Figure S9

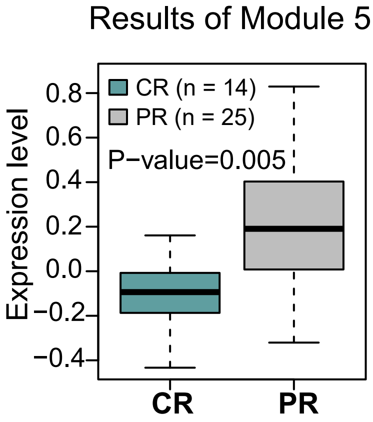
Results of Module 5
CR (n = 14)
PR (n = 25)
P−value=0.005
Expression level
0.8
0.6
0.4
0.2
0.0
−0.2
−0.4
CR
PR

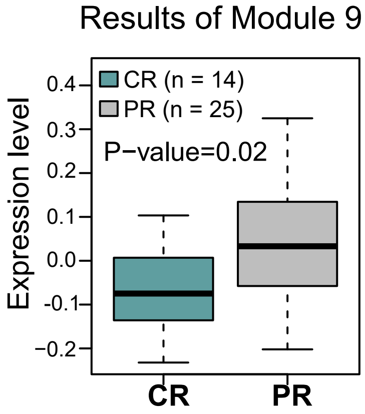
Results of Module 9
CR (n = 14)
PR (n = 25)
P−value=0.02
Expression level
0.4
0.3
0.2
0.1
0.0
-0.1
−0.2
CR
PR
