## Supplementary File for "scLM: automatic detection of consensus gene clusters across multiple single-cell datasets"

**Case study 1: scLM identified tumor-specific modules enriched in specific cell state**

As a case study, we used scLM to analyze our in-house scRNA-seq profiling from 4 NSCLC patients (P1-P4) [36] to identify the co-expressed genes in tumor and normal epithelial cells respectively. In tumor cells (**Figure 5**A, heatmap of latent variables), we discovered 12 co-expressed gene modules in the latent space (T1 – T12). These modules showed clear differences but were consistently concordant across patients (Figure 5A, heatmaps P1 – P4), even though the single cells from different patients presented strong heterogeneity and batch effects (Figure 5B, left panel).

Using the 12 co-expression modules, the single cells were separated into two major clusters. In each cluster, cells from different patients mixed well without interference from batch effects (Figure 5B, right panel), which further support that the co-expression modules are consistent across patients. Interestingly, we found that cluster 1 had higher expression of epithelial functional markers (EMT-related genes) than cluster 2 (Figure 5C). These results indicate that co-expression modules are capable of characterizing specific cell phenotypes.

Similarly, in normal single cells, we observed 13 co-expressed gene modules (N1 – N13) that showed concordant variations in expression levels across individual patients (**Figure S2**). Further comparison of co-expression modules from tumor and normal cells revealed very dissimilar patterns (**Figure S3**A). Some tumor modules (e.g. T11 and T5) had correlations with one normal module, whereas other tumor modules (e.g. T2, T6, and T7) were associated with more than two normal modules. Four tumor modules (T1, T3, T4, and T10) were not correlated with any normal modules, suggesting they were tumor-specific. Meanwhile, we found that these tumor-specific modules had higher expression levels in cluster 1 than in cluster 2 (T10 is shown in Figure S3B; T1, T3, and T4 are shown in **Figure S5**).

To better understand these tumor-specific modules, we examined their enriched biological categories in the REACTOME and KEGG databases (**Figure S6**A-B). T10 was enriched in cell-cell communication, adherence junction, and leukocyte transendothelial migration pathways. Genes in T1 were associated with the circadian clock and the circadian rhythm mammal pathways. T3 was enriched with downregulation of smad2/3/4 transcriptional activity and ubiquitin-mediated proteolysis pathways. No pathways were enriched in the T4 module. Next, we identified the putative upstream transcriptional factor (TF) that mediated these tumor-specific modules (Figure S3D, S6C). Each module was subjected to cis-regulatory motif analysis using the RcisTarget tool to identify their upstream TFs, and only those involved in co-expression modules were retained. TEAD1 and FOXA1 were the major co-expressed regulators of the T10 module. YY1 and ELF2 were regulators of the T1 and T3 modules, respectively, while SRF and POLR2A regulated the T4 module. To further investigate whether these tumor-specific modules have prognostic significance in clinical samples, we used the NSCLC patient data from The Cancer Genome Atlas (TCGA). Remarkably, in patients with Lung Squamous Cell Carcinoma (LUSC), we observed significant associations between overall survival and each tumor-specific module (T10 is shown in Figure S3E, others in **Figure S7**A). However, such association was not seen in patients with lung adenocarcinoma (LUAD) (Figure S7B).

**Case study 2: scLM identified a common program across three types of cancer**

Not only in lung cancer, we also investigated the single cell data from melanoma and HNSCC. For the melanoma dataset, single cells from different patients show strong batch effects (**Figure S8**A, left panel). The co-expressed genes identified by scLM separated the tumor cells into major clusters. In each cluster, cells from different patients mixed well without interference from batch effects (Figure S8A, right panel). For the HNSCC data, the co-expressed genes identified by scLM still achieved to align cells evenly that were not interfered with batch effects from different patients. These results further confirm the consensus properties of the co-expressed genes identified by scLM.

Tumors located in different organs carry different genomic and molecular characteristics but also share common hallmarks that are intrinsic for carcinogenesis. To explore such common hallmarks across different tumor types, we next extended our analysis from NSCLC to the head and neck squamous cell carcinoma (HNSCC) and melanoma, by applying the scLM method to the corresponding public scRNA-seq data. In addition to the 12 co-expression modules in NSCLC, we identified 11 modules in HNSCC and 14 modules in melanoma. To determine the (dis)similarities among the co-expression modules from these three cancer types, we performed a pair-wise comparison of these modules using weighted jaccard similarity, followed by hierarchical clustering. As shown in the diagram (Figure S8B), we found that most branches were dominated by a mixture of cancer types instead of one specific cancer type. For example, one branch contained module 1 from melanoma (melanoma_m1), module 3 from NSCLC (lung_m3), and module 3 from HNSCC (hnscc_m3); whereas another branch contained melanoma_m14, lung_m5, and hnscc_m3. No branch was dominated by a single cancer type, indicating that most modules are not cancer-specific. Importantly, we identified a distinct branch with high similarity among lung_m9, hnscc_m7 and melanoma_m12 modules.

Since the three modules from different cancer types had high similarity, we investigated their potential clinical value further (Figure S8C). In both LUAD and LUSC subtypes of lung cancer, the lung_m9 shows significantly higher expression in tumor than normal samples. In HNSCC samples, the hnscc_m7 is also higher in tumor samples than normal samples. Melanoma module 12 (melanoma_m12) was more pronounced in uveal melanoma (UVM) samples than in skin cutaneous melanoma (SKCM) samples from TCGA. In both HNSCC and uveal melanoma cases, these modules (HNSCC module 7 and melanoma model 12) were associated with poorer survival in patients (with significant p-values of 2.5e-03 and 5.0e-03, respectively). We did not find significant survival difference associated with the corresponding NSCLC module 9 in LUAD and LUSC cases. Overall, these results demonstrate the prevalent malignancy properties of these three similar modules across three cancer types.

These three similar modules substantially overlapped with 91 genes, which were defined as a common program across three cancer types. To gain insights into the biological functions of the common program, we performed enrichment analysis in the Hallmark database (Figure S8D). The MYC targets v1 and hypoxia were the top enriched terms, involving the genes FOS, GAPDH, HLA-A, and NFKB1A. This result suggests the presence of a common intrinsic mechanism of tumor malignancy regardless of cancer types.

With the co-expression modules identified by scLM, it prompted us to test whether the co-expression modules were related with clinical responses to Immune checkpoint inhibitors (ICI). Therefore, we looked into the available RNA-seq cohort collected from 112 melanoma patients prior to ICI treatment (**Figure S9**), to determine the ICI resistance associated co-expression modules in melanoma. Through comparing the post-ICI-complete response (CR) patients to post-ICI-partial response (PR) patients, we found that the melanoma module 5 and module 9 significantly distinguished the ICI-CR from the ICI-PR patients. It suggested that these two modules related with ICI resistance that might provide a predictive signal of ICI therapy response.
